# High-throughput whole-brain mapping of rhesus monkey at micron resolution

**DOI:** 10.1101/2020.09.25.313395

**Authors:** Fang Xu, Yan Shen, Lufeng Ding, Chao-Yu Yang, Heng Tan, Hao Wang, Qingyuan Zhu, Rui Xu, Fengyi Wu, Cheng Xu, Qianwei Li, Peng Su, Li I. Zhang, Hongwei Dong, Robert Desimone, Fuqiang Xu, Xintian Hu, Pak-Ming Lau, Guo-Qiang Bi

**Affiliations:** Center for Integrative Imaging, Hefei National Laboratory for Physical Sciences at the Microscale, and School of Life Sciences, University of Science and Technology of China, Hefei, Anhui, China; CAS Key Laboratory of Brain Connectome and Manipulation, Interdisciplinary Center for Brain Information, The Brain Cognition and Brain Disease Institute, Shenzhen Institutes of Advanced Technology, Chinese Academy of Sciences; Shenzhen-Hong Kong Institute of Brain Science-Shenzhen Fundamental Research Institutions, Shenzhen, Guangdong, China; CAS Key Laboratory of Brain Function and Disease, and School of Life Sciences, University of Science and Technology of China, Hefei, Anhui, China; Key Laboratory of Animal Models and Human Disease Mechanism, Kunming Institute of Zoology, Chinese Academy of Sciences, Kunming, Yunnan, China; Department of Pathology and Pathophysiology, Kunming Medical University, Kunming, Yunnan, China; Institute of Artificial Intelligence, Hefei Comprehensive National Science Center, Hefei, Anhui, China; McGovern Institute for Brain Research, Massachusetts Institute of Technology, Cambridge, Massachusetts, USA; State Key Laboratory of Magnetic Resonance and Atomic and Molecular Physics, Key Laboratory of Magnetic Resonance in Biological Systems, Wuhan Institute of Physics and Mathematics, Chinese Academy of Sciences, Wuhan, Hubei, China; Zilkha Neurogenetic Institute, Center for Neural Circuits & Sensory Processing Disorders, Keck School of Medicine, University of Southern California, Los Angeles, California, USA; USC Stevens Neuroimaging and Informatics Institute, Laboratory of Neuro Imaging (LONI), Keck School of Medicine of University of Southern California, Los Angeles, California, USA; CAS Center for Excellence in Brain Science and Intelligence Technology, Shanghai, China

## Abstract

Whole-brain mesoscale mapping of primates has been hindered by large brain size and the relatively low throughput of available microscopy methods. Here, we present an integrative approach that combines primate-optimized tissue sectioning and clearing with ultrahigh-speed, large-scale, volumetric fluorescence microscopy, capable of completing whole-brain imaging of a rhesus monkey at 1 µm × 1 µm × 2.5 µm voxel resolution within 100 hours. A progressive strategy is developed for high-efficiency, long-range tracing of individual axonal fibers through the dataset of hundreds of terabytes, establishing a “Serial sectioning and clearing, 3-dimensional Microscopy, with semi-Automated Reconstruction and Tracing” (SMART) pipeline. This system supports effective connectome-scale mapping of large primates that reveals distinct features of thalamocortical projections of the rhesus monkey brain at the level of individual axonal fibers.

## Introduction

Given the status of the rhesus macaque (*Macaca mulatta*) as a major experimental animal for modeling human cognitive functions and brain diseases ^1, 2^, a fundamental task in neuroscience and neurology is mapping structural connectivity among different brain regions and neurons (*i*.*e*., the mesoscopic connectome) of the monkey brain ^3, 4^, as those established for the mouse brain ^5, 6^. Connectivity mapping of non-human primate brains has to date relied primarily on bulk labeling of specific brain regions with anterograde and retrograde tracers, followed by interleaved 2D imaging of serial thin sections ^7-9^. However, this approach is tedious and lacks the continuity necessary for tracking individual axons throughout the brain.

Widely used tractography approaches based on diffusion-weighted magnetic resonance imaging (dMRI) are able to image the entire monkey or human brain as a whole, but their anatomical accuracy is inherently limited ^10-12^. Light-sheet microscopy (LSM) combined with whole-brain clearing techniques can image intact mouse brains, but lacks the resolution to distinguish individual axons ^13-17^. Recently developed block-face imaging techniques, including fluorescent micro-optical sectioning tomography (fMOST) ^18^ and serial two-photon (STP) tomography ^19^, have successfully implemented brain-wide axonal tracing in mice, and have opened a new era of connectomic mapping ^20, 21^. However, given that these techniques require several days to image a mouse brain, it is impractical to scale up these methods toward systematic connectomic mapping for macaque or human brains.

To break these technical bottlenecks, we developed an integrative approach consisting of serial sectioning of the brain tissue into thick slices, clearing with primate-optimized uniform clearing solutions, microscopic imaging using ultrafast volumetric on-the-fly scanning, and a semi-automated process for volume reconstruction and axonal tracing. This “SMART” strategy and pipeline (Fig. 1a), owing to ultra-high speed and scalability, overcame several key technical challenges to enable high-resolution mapping of the entire macaque brain. In a proof-of-principle experiment, we generated a projection map from viral-labeled thalamic neurons to the cerebral cortex and unveiled distinct axonal routing patterns in the folded banks along the superior temporal sulcus (*sts*), and carried out efficient semi-automated tracing of neuronal fibers through the near petabyte dataset of the monkey brain.

**Fig. 1.**
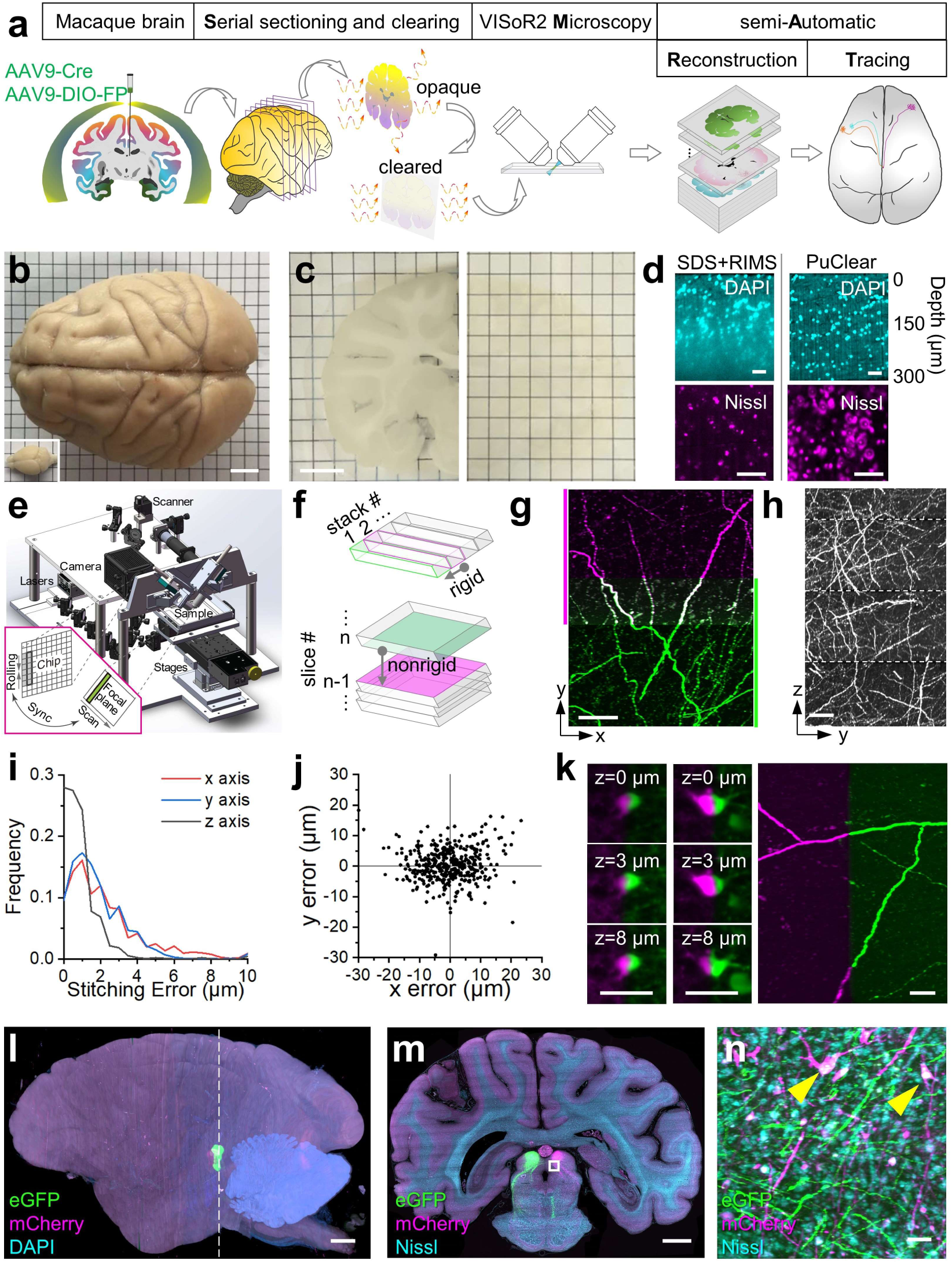
The SMART approach for high-throughput mapping of a rhesus macaque brain at micron resolution. (**a**) The SMART pipeline. (**b**) A macaque brain compared with a mouse brain (bottom left). (**c**) A macaque brain slice before (left) and after (right; flipped) PuClear treatment. (**d**) Comparison between SDS-and RIMS-based clearing, and PuClear treatment. Top panels: WM areas stained with DAPI. Bottom panels: cortical areas stained with a fluorescent Nissl dye, NT640. (**e**) The VISoR2 imaging system. Inset: a schematic showing the synchronization mechanism between laser scanning and camera readout. (**f**) A schematic for volume stitching within a single slice (top) and between slices (bottom). (**g**) Maximum intensity projection (MIP) image of a 100-µm coronal section of two adjacent stacks (separately color-coded) with merged overlapped regions after stitching. (**h**) MIP of a 50-µm virtual section from 4 consecutive slices. (**i**-**j**) Distributions of the errors of intra-slice (i) and inter-slice (j) stitching. (**k**) Stitched neurons (each shown in 3 z-sections) and axonal branches cut into two adjacent brain slices (separately color-coded). (**l**) A reconstructed macaque brain with viral labeling of bilateral SC areas. (**m**) MIP of a 30-µm coronal section indicated with a dashed line in (l). (**n**) Magnified view of the boxed region in (m). Arrowheads indicate neurons co-labeled by the virus and fluorescent Nissl staining. Scale bars: (b), (c), 10 mm; (d), (k) and (n), 50 μm; (g-h), 100 μm; (l) and (m), 5 mm.

## Results

### Serial sectioning, clearing and high-throughput imaging of the macaque brain

The first major challenge for imaging large brains is sample preparation. The difficulty of reagent penetration increases exponentially with tissue thickness, making it exceedingly difficult to achieve uniform histological staining or clearing of the whole monkey brain, which is more than 200 times larger than a mouse brain (Fig. 1b) ^8^. We therefore chose to section the brain into slices before subsequent clearing and imaging. A robust workflow was established with hydrogel-based embedding to minimize tissue loss and distortion during sectioning and clearing (Supplementary Figs. 1 and 2; online Methods). A macaque brain was sectioned into about 250 consecutive 300-µm slices which were treated with a <u>p</u>rimate-optimized <u>u</u>niform <u>clear</u>ing method (PuClear) that combines Triton X-100-based gentle membrane permeabilization with high refractive index matching (Fig. 1c). Unlike the widely used sodium dodecyl sulfate (SDS)-based clearing methods such as CLARITY ^13, 14^ that we found inadequate for imaging clearly through the white matter (WM) of primate tissue, PuClear has a refractive index of 1.52, yielding uniform transparency through the full depth of 300-µm slices including WM areas (Fig. 1d, top panels). Importantly, PuClear preserves the morphology of neurons labeled by Nissl stain (Fig. 1d, bottom panels) and showed excellent compatibility for both immunostaining and retrograde tracing using cholera toxin subunit B (CTB) (Supplementary Figs. 3 and 4).

Uniform clearing of thick brain slices also allowed us to overcome a second major challenge, the long duration of time required for imaging a large brain at high resolution. For this, we developed a new iteration of our recently reported synchronized on-the-fly-scan and readout (VISoR) technique ^22^. This improved “VISoR2” system is optimized for ultrahigh-speed volumetric imaging of the larger monkey brain sections (Fig. 1e). Besides instrumental upgrades including long-travel linear stages and a more compact and stable light-path, the new system was implemented with an optimized control sequence for the sCMOS camera, the illumination laser, and the galvanometer scanner (Supplementary Fig. 5; online Methods).

We achieved 250 Hz blur-free imaging of a 0.7×2 mm^2^ field of view containing the optical section of the slice with smooth stage movement at any speed ranging from 0.5 to 20 mm/s. This configuration corresponds to a voxel resolution of 1.0×1.0×(1.4∼56) µm^3^ and a continuous data rate of 400 million voxels per second (Supplementary Fig. 5; Supplementary Video 1). Thus the system is capable of imaging a mouse brain that is serially sectioned, cleared and mounted on a single glass slide within 30 min at 1.0 µm × 1.0 µm × 2.5 µm resolution. Consequently, the collection of ∼80 million single-channel images (2048 × 788 pixels each) for all slices from one macaque brain only took 94 hours imaging time. This VISoR2-based imaging across three channels resulted in 750 terabytes of data for a rhesus macaque brain that was labeled via co-injection of AAV cocktails mixed by an adeno-associated virus (AAV) carrying Cre recombinase and another AAV carrying Cre-dependent fluorescent proteins (FPs) reporter, either eGFP or mCherry, into the left and right superior colliculus (SC), respectively.

### Reconstruction of the entire macaque brain

While the VISoR2 microscopy with significantly improved imaging speed overcame the second challenge in primate brain mapping, it generated a third challenge: the analysis of such large dataset. Available tools have been effective in handling terabyte-level multi-tile images, but these tools cannot be used for non-overlapped image tiles ^23, 24^, or lack automation for multi-hundred terabyte data ^25^. We therefore developed a custom software tool that implements automated volume stitching (Fig. 1f, Supplementary Fig. 6), including rigid-transformation-based 3D intra-slice stitching (Fig. 1g) and non-rigid-transformation-based inter-slice alignment (Fig. 1h, Supplementary Video 2). Attesting the strong performance of this tool, we found that the intra-slice stitching errors were ∼2 µm for each axis (Fig. 1i) and the inter-slice alignment error of AAV labeled axons was ∼8 µm (Fig. 1j). Importantly, tissue loss between consecutively sectioned slices was insignificant, as seen in the precise alignment of truncated cell bodies and axonal branches (Fig. 1k). As we demonstrate below, this precision was sufficient for visually tracing axonal projections. In practice, we reconstructed the whole brain at a coarse voxel resolution (10 × 10 × 10 µm^3^) to obtain an overview of brain structures, and also established a robust transformation framework for on-demand reconstruction of user-specified regions of interest (ROIs) at full resolution for detailed analysis of axonal projections (Fig. 1l-n, and Supplementary Fig. 7).

### Mesoscopic mapping of thalamocortical projection

To demonstrate the capacity of our system for mesoscopic mapping of neuronal projections across the entire monkey brain, we bilaterally injected AAV cocktails into the left and right medial dorsal nucleus of the thalamus (MD), with minor leakage to nearby areas (Supplementary Fig. 8). The MD is known to generate dense projections to the prefrontal cortex (PFC) ^26, 27^. VISoR2 imaging and 3D reconstruction of this monkey brain allowed for visualization of the global distribution of axons originating from the injection sites and projecting to the cerebral cortex (Fig. 2a). From the 3D volume and a series of virtual sections, it was clear that bundled fibers from the injection sites traveled through the internal capsule in horizontal, obliquely lateral, and upward directions (Fig. 2b, Supplementary Video 3), before continuing on towards the frontal lobe, where they primarily targeted the posterior orbitofrontal area (OFC) (Fig. 2c, Supplementary Fig. 9), largely consistent with previous reports of the MD projections ^26, 28^. Furthermore, the resolution of the original images was sufficient for us to visualize individual axons and branches, and to distinguish whether the fibers were passing by or making terminal arborizations. For instance, our PFC mapping revealed that labeled axons terminate in layer III and layer IV (Fig. 2d).Besides the canonical target areas of the MD in the ipsilateral PFC with high-density arborizations of labeled axons, we also observed various lower density, yet significant, fiber branches and arborizations in other areas such as the ipsilateral secondary somatosensory cortex (SII) (Fig. 2e,f), which is known to be a target of the ventral posterior inferior nucleus (VPI) and ventral posterolateral nucleus (VPL) ^29^, and the temporal lobe, including superior temporal sulcus (*sts*) dorsal (TPO) and ventral (TEa) bank areas (Fig. 2e,g-h, Supplementary Fig. 9). All projection targets observed in this animal are summarized in Supplementary Table 1 and visualized in Fig. 2i in a cortical map after flattening (Supplementary Fig. 10). Thus, the high resolution and sensitivity of SMART imaging may help reveal previously unidentified cortical targets of the MD, although some of these observations could be due to inadvertent labeling of neurons in other nuclei close to the injection sites. More precise injection or a sparse labeling approach that allows complete single neuron tracing will help resolve such uncertainties and unveil detailed thalamocortical projection maps.

**Fig. 2.**
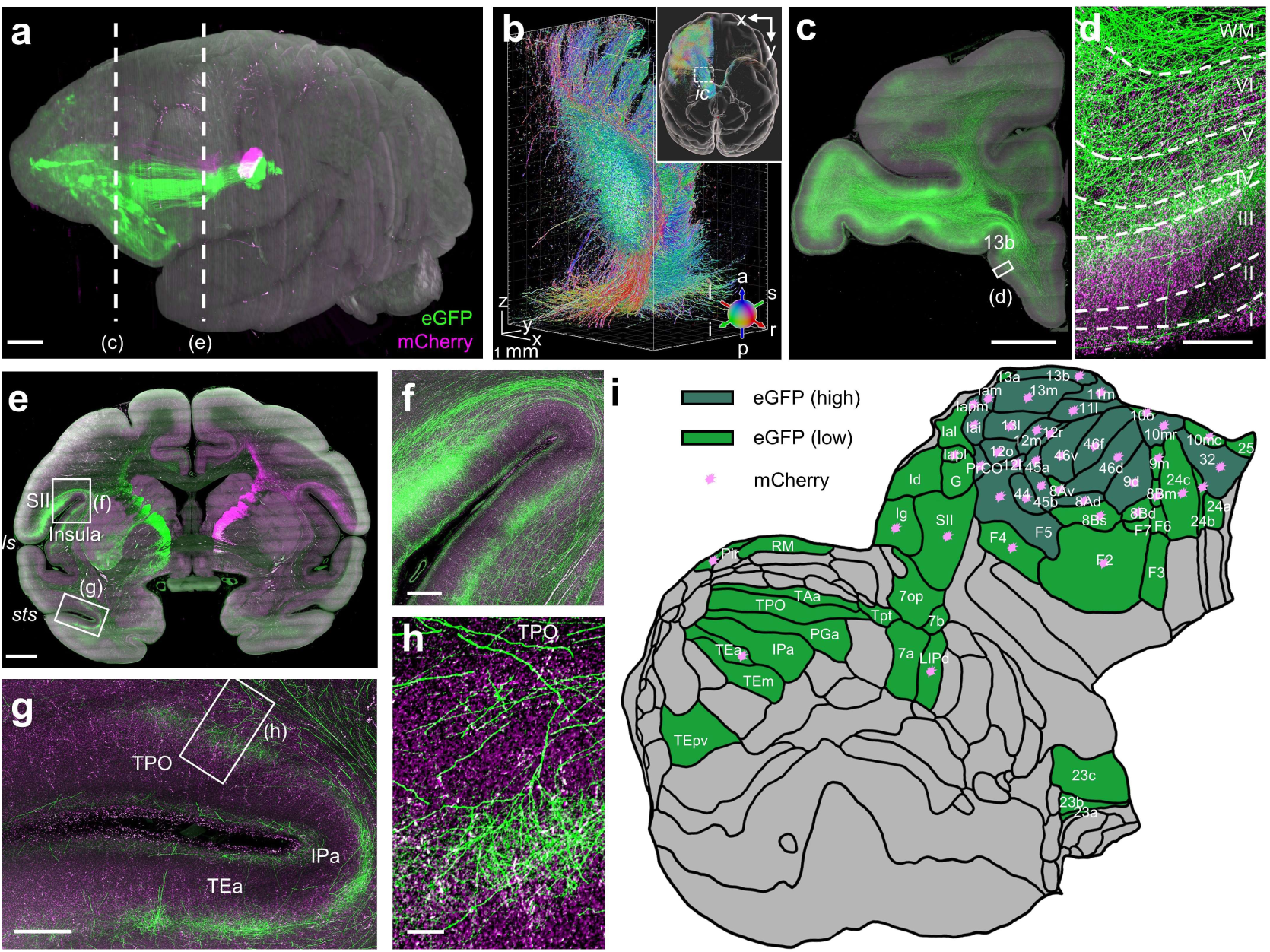
Mesoscopic mapping of the MD projection. (**a**) A reconstructed macaque brain with viral labeling of bilateral MD areas. (**b**) Fiber orientation image of the brain (inset) and a magnified view of the left *ic* (box region). Only the eGFP channel is displayed, where red, green, and blue colors represent the right/left, anterior/posterior, and superior/inferior orientations, respectively. (**c-h**) MIPs of two 300-μm coronal sections (c, e), and the magnified views (d, f, g, h) of the boxed regions specified in (c), (e), and (g). Cortical layers were drawn based on the autofluorescence patterns in each channel. (**i**) Summarized cortical flat map showing high-and low-density distribution of axonal projections to the ipsilateral hemisphere from the injection sites revealed by eGFP and mCherry expression. Scale bars: (a), (c), (e), 5 mm; (d), (g), 500 μm; (f), 1 mm; (h), 100 μm. Green codes the eGFP channel and magenta codes the mCherry channel in all panels except in (b). Acronyms: *ic*, internal capsule; *ls*, lateral sulcus; *sts*, superior temporal sulcus; others are listed in Supplementary Table 1.

### SMART reveals distinct axonal routing strategies in local cortical folds

The resolution of our system also allowed for identification of fine features of individual axons in the projection sites where the fibers were not excessively dense. As an example, we reconstructed a full-resolution volume of the areas near the *sts* and traced 30 randomly selected axonal segments (Fig. 3a,b and Supplementary Video 4) using custom tracing software. Intriguingly, in these folded areas, the axons projecting to layers III/IV of the TEa typically navigated to the dorsal side first, before separating into two major groups: one group made sharp turns in the WM (Fig. 3c, top panel), traveling along the boundary of the WM (Fig. 3c, bottom panel); the other group made right-angle turns and traveled through the superficial cortical layers (Fig. 3d). We observed 4 distinct classes of turning patterns for these axons (Fig. 3e-h). Such differences in the micro-organization of these afferent axons may underlie their functional diversity, especially when they make putative *en route* connections with different sets of neurons positioned within various local circuits.

**Fig. 3.**
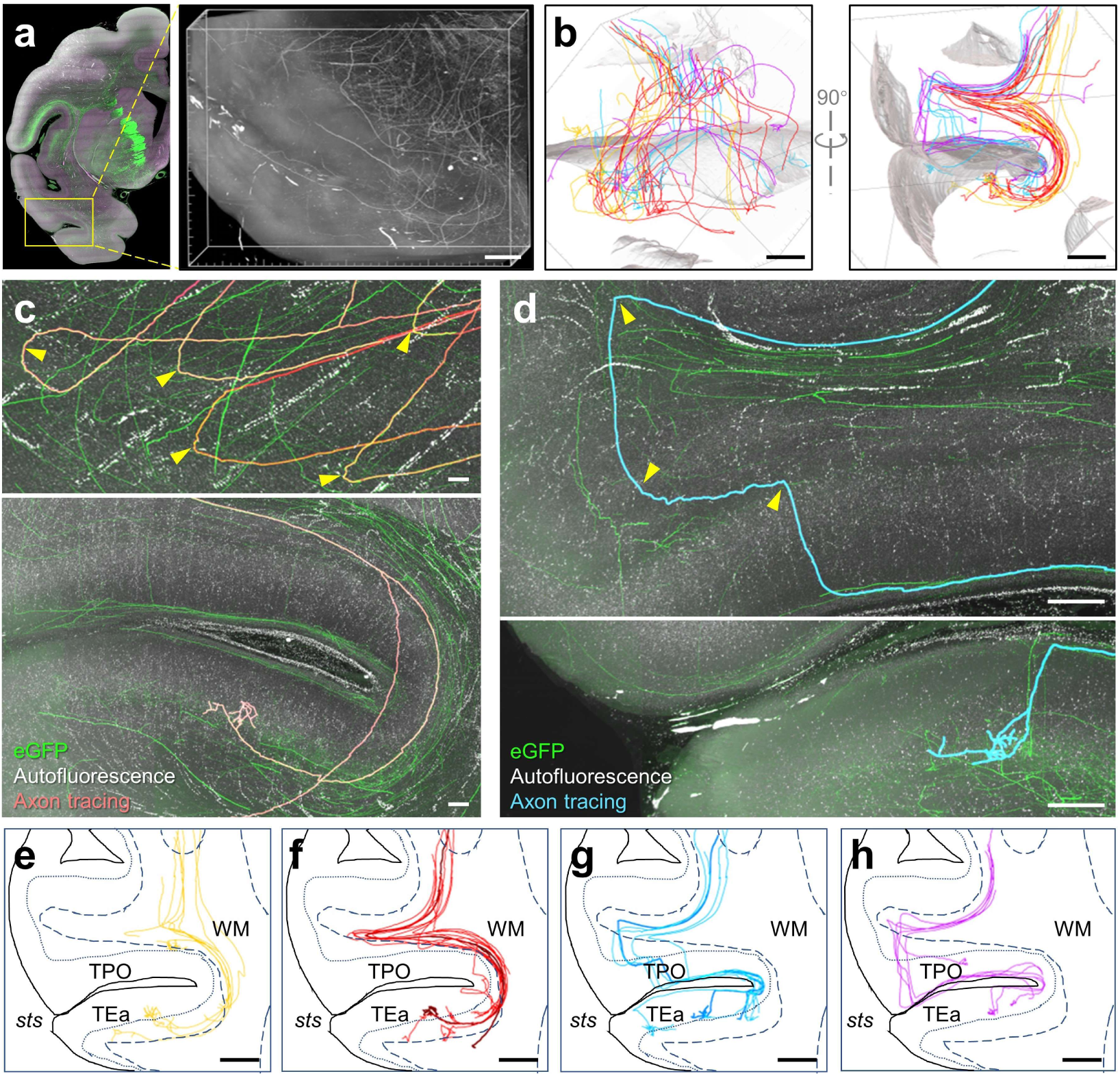
Organization of axonal fibers in cortical folds. (**a**) Overview of a 16×11×11 mm^3^ image volume surrounding the *sts* from an MD-injected macaque. (**b**) 30 traced axonal segments shown from two perspectives. Curved brain surfaces were segmented and rendered as a shaded reference to help visualize the location of axon trajectories. (**c**) Example images showing the sharp turns (arrowheads) of 5 axons (top) in the white matter (WM) and the arbor area of an axon (bottom). (**d**) Example images showing the right-angle turns (arrowheads) of an axon (top) and its arbors (bottom). Images in the top panels of (c) and (d) are MIPs of 450-µm virtual sections resliced in an orientation parallel to the fibers. The bottom images are MIPs of 1200-µm sections. (**e-h**) Four axonal navigation patterns, including: those following short paths (e), those making sharp turns (f), those making two-step right-angle turns (g), and those making right-angle turns (h). The boundaries of cortical surfaces are illustrated as solid lines; boundaries of the WM are illustrated as dashed lines. Cortical layer IV can be recognized as a dim band in the autofluorescence images; it is drawn with dotted lines. Fibers shown in (c) and (d) are highlighted in (f) and (g), respectively. Scale bars: (a-b), (e-h), 2 mm; (c) and (d), 200 μm. Acronyms: *sts*, superior temporal sulcus; TPO, *sts* dorsal bank area; TEa, *sts* ventral bank area; WM, white matter.

### Brain-wide axonal tracing of macaque neurons

With the capacity of resolving individual axons, we set out to trace the long-range, whole-brain projections of macaque axons but encountered yet another challenge. Whereas convenient tools have been developed for brain-wide tracing of individual axons in mouse ^20, 30-32^, it is computationally challenging to scale up these tools to handle the whole brain volume of a macaque at full resolution. In addition, the reconstructed local image is often of somewhat lower quality than the raw image, partly because of errors introduced by non-linear deformation and interpolation steps that are implemented to achieve global consistency. Therefore, we developed a computationally efficient strategy for progressively tracing axons in blocks directly from raw images. For this, we first generated a relatively low resolution (10 × 10 × 10 μm^3^) reconstruction volume of the macaque brain, and established a “SMART positioning system” (SPS) that employs a set of bidirectional transformations to enable mapping between the initially defined whole-brain coordinate system and the corresponding data from each raw image (Fig. 4a).

**Fig. 4.**
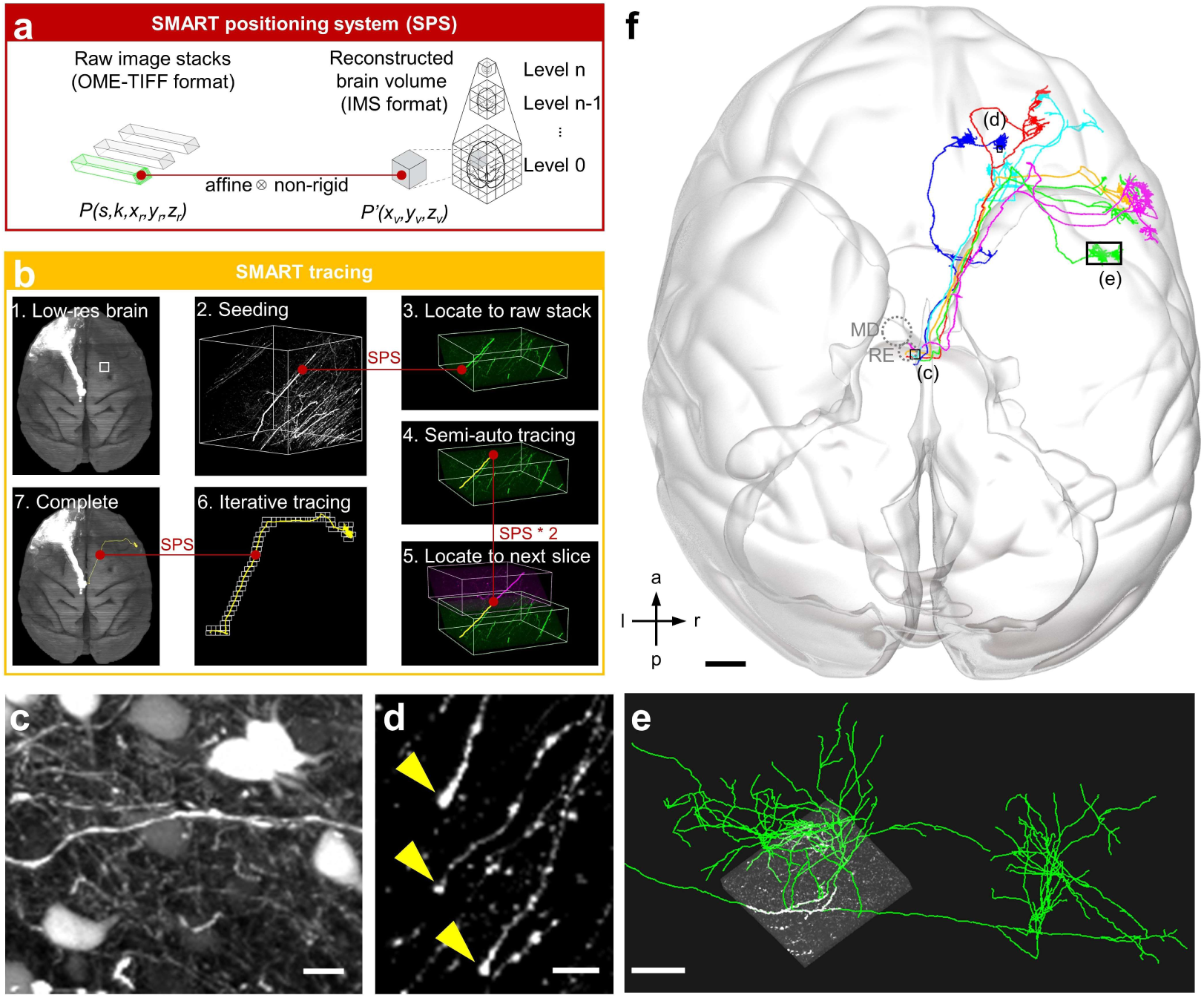
Brain-wide tracing of axonal projections. **(a)** The SMART positioning system (SPS) maps between any position (*x*_*r*_, *y*_*r*_, *z*_*r*_) in the raw image stack (*k*) of a brain slice (*s*) and the corresponding point (*x*_*v*_, *y*_*v*_, *z*_*v*_) in the space of the reconstructed brain volume. (**b**) Step-by-step workflow for SMART-based axonal tracing. Starting from a low-resolution image (b1), a seed node was selected (b2) and located via SPS in the raw image stack (b3); subsequent semi-automated tracing was conducted (b4) to navigate through to the edge of the slice and beyond into the neighboring zone of the adjacent slice as identified by SPS (b5). The entire axon can then be traced by iteratively applying steps 4 and 5 (b6), and finally visualized in the reconstructed brain space by reverse transformation of traced coordinates via SPS (b7). (**c-d**) Tracing was terminated at the injection site where the axons were too dense (c) or at axonal termini (d). (**e**) Magnified view of axonal terminals overlapped with a raw image block. (**f**) Six thalamocortical axons projecting to the right hemisphere from the left MD or RE areas are shown. Scale bars: (c), 20 μm; (d), 10 μm; (e), 500 μm; (f), 5 mm. Acronyms: MD, mediodorsal nucleus; RE, reunions nucleus.

An iterative workflow was then established to trace the relatively sparsely labeled axons projecting to the contralateral hemisphere (Fig. 4b). In this scheme, bright fiber trunks were first identified in the low-resolution whole-brain image, followed by semi-automatic tracing in full-resolution raw image blocks that were dynamically loaded as tracing progressed along the axon track; this process proceeded until reaching the injection site (where the fibers were too dense) (Fig. 4c) or reaching the nerve ending of each axonal branch (Fig. 4d,e). When necessary, any misalignment between adjacent slices from errors in the automatic registration process was manually corrected based on the continuity of foreground neuronal fibers and background microvasculature (Supplementary Fig. 11).

Using this progressive SMART tracing strategy, we tracked all 28 randomly selected bright fiber trunks retrogradely back into the injection site and anterogradely to their branching points. These fibers travel in parallel in a bundle within the internal capsule before branching out into divergent cortical areas (Supplementary Fig. 12; distances before branching: 26.8±1.6 mm; n=28). We also selected 6 from those fiber trunks and mapped out their full terminal arborizations (Fig. 4f and Supplementary Video 5). For tracing each axon, only 1.7±0.5 % of raw images were sequentially accessed (total size of raw images: 238 TB; size of images accessed during tracing: 4.1±1.2 TB; n=6), with multiple 600 MB image blocks (2048×788×200 voxels each) loaded into the memory at each time, a workload manageable by a personal computer. Notably, most of these axonal fibers form clustered arborizations in confined cortical regions, with negligible subcortical arborizations (3.4±2.6 % of total axonal length; n=6), in striking contrast to the previously mapped mouse thalamic projections from the MD and nearby reunions nucleus (RE) (Supplementary Fig. 13; Supplementary Table 2).

## Discussion

Owing to its implementation of optimized tissue slicing and clearing, ultrahigh-speed imaging techniques, and efficient analysis tools for processing near-petabyte-scale datasets, SMART bridges the gap in our understanding of functionally impactful differences between rodent and human brain architectures, specifically by enabling the efficient mapping of primate brains at subcellular resolution and supporting brain-wide, long-range tracing of individual axons. Indeed, our proof-of-concept study has already begun to reveal potential new targets of primate thalamocortical projections and to highlight distinct properties of individual axons, including their long trunks and striking turning patterns as they progress towards cortical targets. Although this initial study only allowed for tracing of a small number of single fibers because of the very dense labeling and relatively low throughput of semi-automatic axonal tracing, much sparser labeling is achievable by lowering the concentration of Cre recombinase-carrying AAV in the viral injection cocktail ^33, 34^, and combining with high-performance computing (HPC) and automated tracing techniques, it is expected that the SMART system will allow for mapping the full morphology of a potentially huge number of individual neurons ^20, 30, 35^, thus paving the way toward a truly connectome-scale understanding of the primate brain.

It should also be noted that SMART is compatible with widely used experimental techniques for histological labeling, thereby supporting analysis of samples not amenable to viral labeling, for example postmortem human brains ^36^. The strategies underlying SMART, including non-overlapped physical slicing and computational stitching, high-throughput blur-free imaging, and progressive tracing in the raw image stacks, are all readily scalable and applicable to other biological samples, including internal organs and even the whole bodies of various species that labeled with antero-or retrograde transneuronal transporting viruses ^37^. Application of these techniques have the potential to yield unprecedented understanding of brain architecture, and high-precision, systems-level insights about the development, basic functions, and neurological pathology of the entire nervous system.

## Methods

### Labeling viruses

For anterograde neural labeling, recombinant adeno-associated viruses (rAAVs) were generated by transient triple transfection of HEK293 cells as previously reported ^38^. Cap serotype 9 was chosen to package the AAV vectors to achieve high transduction levels and high titers (>10^12^ vg/mL). A strong promoter CAG and transcription control element WPRE were chosen to construct pAAV-CAG-Dio-EGFP-WPRE-pA or pAAV-CAG-Dio-mCherry-WPRE-pA constructs for stable fluorescent protein (FP) expression in primates ^39^. To increase neuronal specificity, we used the hSyn promoter to construct pAAV-hSyn-Cre-WPRE-pA to serve as a controller of FP-expressing vectors.

### Mice

Eight-week-old male C57BL/6 and Thy1-YFP-H (Jax: 003782) mice were used in this study for prototyping the sample preparation and imaging methods. All mice experiments were carried out following protocols approved by the Institutional Animal Care and Use Committees of the University of Science and Technology of China (USTC). All mice used in this study were group-housed with a 12-hour light/dark cycle (lights on at 7 a.m.) with free access to food and water.

4% hydrogel monomer solution (HMS) was prepared for perfusion by mixing 40% (w/v) acrylamide (4% final concentration; V900845, Sigma), 2% (w/v) bisacrylamide (0.05% final concentration; V3141, Promega), 10× phosphate buffer saline (PBS; 1× final concentration; 70011044, ThermoFisher), 8% (w/v) paraformaldehyde (PFA; 4% final concentration; 157-8, Electron Microscopy Sciences), distilled water, and VA-044 thermal initiator (0.25% final concentration; 223-02112, Wako) ^13^. The perfusion procedures were carried out following a modified protocol based on a previous study ^22^. Mice were deeply anesthetized with 1% sodium pentobarbital solution, followed by transcardial perfusion with 20 mL of 37°C PBS, 20 mL ice-cold PBS, and finally 20 mL of ice-cold 4% HMS. All solutions were perfused at a uniform rate of 10 mL/min. Mouse brains were harvested and immediately placed into 20 mL ice-cold 4% HMS and incubated at 4°C for 24-48 h to allow further diffusion of the hydrogel monomers into the tissue.

### Monkeys

Three adult, 10-year-old, male rhesus macaques (*Macaca mulatta*) were used in this study. Macaques were obtained from the breeding colonies of the Primate Research Center of Kunming Institute of Zoology, Chinese Academy of Sciences (KIZ, CAS), which was accredited by the Association for Assessment and Accreditation of Laboratory Animal Care (AAALAC International). The experimental procedures were approved by the Institutional Animal Care and Use Committee (IACUC) of KIZ, CAS (IACUC No. IACUC18018).

In order to prevent gastric regurgitation caused by the anesthesia, fasting and water deprivation were implemented for at least 6 h before surgery. Animals were anesthetized using 10 mg/kg ketamine hydrochloride injection (i.m., 50 mg/mL, Zhong Mu Bei Kang pharmaceutical industry limited company, China) and maintained with 20 mg/kg pentobarbital sodium (i.m., 40 mg/mL, Merck, Germany). During the surgery, heart rate and core temperature were monitored using a rectal probe and ECG monitor. MRI assisted brain region positioning was used for accurate encephalic injection as previously described ^40^. MRI scanning was performed using a 3-T scanner (uMR770, United Imaging, China) with a 12-channel knee coil. FP-expressing rAAVs (pAAV-CAG-Dio-EGFP-WPRE-pA, 4.70×10^12^ vg/mL, 2 μL, for the left hemispheres, or pAAV-CAG-Dio-mCherry-WPRE-pA, 3.15×10^12^ vg/mL, 2 μL, for the right hemispheres) and Cre-expressing rAAVs (pAAV-hSyn-Cre-WPRE-pA, 2.09×10^12^ vg/mL, 2 μL) were mixed at a 1:1 ratio for each injection. In this study, a macaque was injected with these AAV cocktails at both sides of the superior colliculus (SC, AP:-4 mm; ML: ±3 mm; DV:-38 mm) and another macaque was injected with these mixtures at both sides of the mediodorsal nucleus (MD, AP: 1 mm; ML: ±3 mm; DV: -38 mm). A third monkey was injected with a cholera toxin subunit B-Alexa Fluor 647 conjugate (CTB-AF647; C34778, Invitrogen) at the quadrigeminal cistern. The duration of injection was more than 20 minutes (including 5 minutes each after inserting and before withdrawing the microsyringe). Antibiotics were used for 3 d after surgery. Injection sites were further confirmed from reconstructed whole-brain images.

Transcardial perfusions were carried out 8 weeks after surgery. Thirty min after anesthesia, each animal was sequentially perfused with the following solutions at the specified speed: PBS 8 L (37°C, 10 mL/s), PBS 1 L (4°C, 1.5 mL/s), 4% HMS 1L (4°C, 1.5 mL/s), and 4% HMS 1 L (4°C, 0.3 mL/s). The brains were extracted immediately after perfusion within 30 min.

### Tissue embedding and slicing

A post-fixation procedure with hydrogel was set up for crosslinking proteins and minimizing tissue loss. Immediately after the macaque brain was excised, it was immersed into 500 mL of 4% HMS and stored at 4°C for 1 week before embedding to allow penetration of the fixatives. Then the brain was immersed in the embedding solution, a 1:1 mixture of 4% HMS (2% final concentration) and 20% bovine serum albumin (BSA; 10% final concentration; V900933, Sigma), incubated at 4°C for 1 week, polymerized at 37°C for 4-5 h, and washed 3 times in PBS to remove residual reagents. Embedding with the mixture of HMS and BSA provides not only *in situ* fixation of proteins^13^, but also high material stiffness and toughness for preserving slice integrity during sectioning (Supplementary Fig. 2). Embedded brains were sectioned into about 250 pieces of 300-µm-thick slices using a vibroslicer (Compresstome VF-800, Precisionary Instruments). All slices of each brain were collected and each slice was placed in a Petri dish with 40 mL PBS and stored at 4°C.

### Sample clearing

The PuClear clearing method was established based on previously reported CLARITY ^13^, CUBIC ^17^ techniques with optimization for primate brain tissues, consisting of membrane permeabilization and refractive index (RI) matching. Brain slices were first treated with a high concentration Triton X-100 solution (5% in PBS; T928, Sigma) for 3-4 d at 37°C with gentle shaking to adequately increase membrane permeability, and were then washed with PBS 3 times. High refractive index (RI) solution was prepared by mixing 50 wt% iohexol (29242990.99, Hisyn Pharmaceutical), 23 wt% urea (A600148-0002, Sangon), 11 wt% 2,2’,2”,-nitrilotriethanol (V900257, Sigma), and 16 wt% distilled water. The final refractive index of PuClear RI-matching solution is 1.52. Before imaging, brain slices mounted onto the glass substrates were incubated in this solution for at least 1 h to allow the sample getting optically transparent.

### Sample staining

Staining was performed after PuClear perforation and before mounting. For immuno-labeling, membrane-perforated slices were placed into Petri dishes and immersed in blocking solution (5% (w/v) BSA in 0.3% PBST) overnight. After that, samples were incubated with the primary antibody in 0.3% PBST for 3-4 d followed by 3 washes with PBS. Subsequently, samples were incubated with the secondary antibody in 0.3% PBST for 2-3 d, followed by 3 times washing with PBS. Dishes were kept at 4°C during blocking, staining, and washing with gentle shaking. For fluorescent Nissl staining, the blocking step was skipped and the slices were incubated with NT640 in 0.3% PBST for 3-4 d at 37°C followed by washing 3 times with PBS. For DAPI staining, samples were incubated with DAPI stock solution for 1 d at 37°C followed by washing 3 times with PBS.

The following antibodies and dyes were used in this study (name, company, catalog number, dilution): Polyclonal Rabbit Anti-Glial Fibrillary Acidic Protein (GFAP), Dako, Z0334, 1:100; Anti-Tyrosince Hydroxylase Antibody, Milipore, MAB318, 1:500; Alexa Fluor 647 AffiniPure Donkey Anti-Mouse IgG (H+L), Jackson ImmunoResearch Laboratories, 715-605-151, 1:200; Alexa Fluor 488 AffiniPure Donkey Anti-Rabbit IgG (H+L), Jackson ImmunoResearch Laboratories, 711-545-152, 1:200; NeuroTrace 640/660 Deep-Red Fluorescent Nissl Stain (NT640), ThermoFisher, N21483, 1:200; DAPI, Beyotime, C1006, stock solution.

### VISoR2 microscope

We designed and built the VISoR2 microscope based on the VISoR technique described previously for mouse brain imaging ^22^, with long-range sample stages and major upgrades for improving its stability, repeatability, as well as the practical imaging speed. The microscope was equipped with four lasers with wavelengths of 405 nm, 488 nm, 552 nm, and 647 nm (all from Coherent). Incident light was combined and illuminated onto a galvo scanner (GVS011, Thorlabs). The position of the scanner was in conjugation with the back focal plane of an illumination objective (10X/NA 0.3, Olympus) via two coupled relay lenses (F=150mm, Thorlabs). 1-D scanning of the scanner generated an illuminating plane that was overlapped with the focal plane of an imaging objective (10X/NA 0.3 or 20X/NA 0.5, both from Olympus). Emission light was filtered with bandpass filters (450/50, 520/40, 600/50, and 700/50 for the four laser sources, respectively; all from Semrock). Images were collected on a CMOS camera (Flash 4.0 v3, Hamamatsu) through a tube lens (IX2-TLU, Olympus) and a 0.63X adapter (TV0.63, Olympus). The objectives were both positioned at a 45-degree incline to the samples. A linear stage (DDSM100, Thorlabs) and a stepper stage (LTS150, Thorlabs) were used for X-and Y-axis movements, respectively, and a stepper stage (MCZ20, Zaber) was used for Z movement.

The devices were controlled by custom software written in C++. To maximize the imaging throughput, the camera was aligned in the center of the light path and its readout was bi-directional in an “*external trigger syncreadout*” mode. During imaging, the X stage moved smoothly at a speed ranging from 0.5 mm/s to 20 mm/s for different resolution requirements. To avoid blurring due to motion, we synchronized the lasers, the scanner, the camera, and the stages with a DAQ board (NI PCIe-6374, National Instruments). The Y-step size was set to include 10% overlapped regions between adjacent image stacks.

Each time the X-stage finished accelerating and moved at a constant speed, it generated a rising edge signal which triggered a timer in the DAQ board to start signaling, continuously generating signals for lasers, the scanner, and the camera until imaging was complete, resulting a stack of image frames of the 45-degree oblique optical sections of the sample. 400 million voxels were acquired per second, approaching the maximum data rate of the CMOS camera. Multi-color imaging was implemented by sequentially imaging individual channels and computationally registering all channels after all images were acquired.

### Workstation

The image acquisition software was run on a workstation equipped with an Intel Xeon E5-2680 CPU, an NVIDIA GTX 1060 graphics card, and 128 GB memory. It was equipped with 2 disk arrays each consisting of 8 SSDs configured in RAID5 for alternatively acquiring and transporting data to a remote petabyte storage server connected via 10 Gbps fiber-optic intranet. Windows 10 Pro 64-bit operating system was run on this workstation.

### Data Management and Compression

The VISoR2 system acquired about 20,000 image frames for a typical monkey brain slice, consisting of several image stacks, each generated during a one-way uniform motion of the X stage. We adopted the BigTIFF format instead of standard TIFF to allow storing all images of a slice into a single file sized ∼200 GB with embedded OME-XML metadata, adapted to the Bio-Formats plugin in ImageJ/Fiji ^41, 42^. We also forked this plugin to provide additional visualization features optimized to these large OME-TIFF images (https://github.com/dinglufe/bioformats). A single-channel VISoR2 image volume of a whole rhesus monkey brain contained ∼1×10^14^ voxels of 16-bit depth which occupies ∼250 TB of storage. An efficient image compression method is necessary for handling such a large dataset, which requires not only a high compression ratio but also low computation resource consumption to compromise the heavy data load. We used the lossless Lempel–Ziv–Welch (LZW) algorithm to compress the raw images ^43, 44^, which typically reached a compression ratio of 2:1. Furthermore, we also used a slightly lossy compression strategy by truncating the 4 least significant bits and performing a proper rounding to the higher 12 bits of each 16-bit voxel value of the original images, followed by additional LZW compression. This method practically reaches a compression ratio of 8:1 with little computation time and CPU usage and worked in real-time during image acquisition. The theoretical peak-signal-noise-ratio (PSNR) of this method achieves 83.0, as determined by 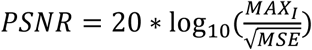, where *MAX*_*I*_ equals 65535, the maximum possible voxel value of a 16-bit image, and MSE is the rounding error, which was determined by 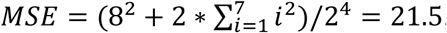, supposing the values of the lower 4 bits follow a uniform distribution. More importantly, this method is seamlessly compatible with all visualization and analysis tools developed for TIFF images.

### Automatic whole brain reconstruction

The volume reconstruction was automatically performed by custom software written in Python.

#### Intra-slice stitching

Raw images of a brain slice were organized as a set of image stacks, with metadata of their physical positions recorded from the output of the X-, Y-, and Z-stages. Then raw image substacks from the overlapped region of two adjacent image stacks, each consisting of 100 continuous images, were sampled to calculate the stitching translation between these two stacks. The stitching translation was determined as the shift that minimized their normalized cross-correlation (NCC) in virtue of the open-source tool *elastix* (http://elastix.isi.uu.nl). The stitched image stacks were then generated by resampling raw images.

The precision of this intra-slice stitching method was calculated by evaluating the NCC between small random ROIs cut from the overlapped regions of two stitched stacks, with one stack fixed and the other moving in a 20 px × 20 px × 20 px window. The stitching error was determined as the shift that minimized the NCC and was further refined through quadratic interpolation of three points nearest to the NCC minimum and then taking the sub-pixel location corresponding to the minimum of the quadratic curve. The sub-pixel refinement formula is:

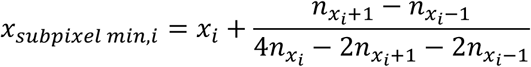

where *x*_i_ is the i-th axis position of the minimal value in the NCC array, and 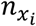 is the NCC value at the position *x_i_*.

#### Channel alignment

For calculating the precise displacement of each stack among sequential multi-channel imaging, one of the channels (usually the eGFP channel) was chosen as the reference channel and each other channel was aligned to the reference channel stack by stack. Several pairs of image substacks consisting of 100 continuous frames were sampled from both this channel and the reference channel at the edges of the brain slices detected by a brightness threshold. The contours of the brain slices provided autofluorescence features for computational alignment. The images were filtered using a gradient magnitude filter, then the translation between each pair of image substacks was calculated using the mutual information metric with *elastix*. The median value of the translation calculated from all the pairs of image substacks was determined as the displacement between this channel and the reference channel.

#### Flattening

The upper and lower surfaces of the brain slice in 3D images were identified and digitally flattened in this step. The upper and lower surfaces were represented by two images *H*_*U*_ and *H*_*L*_. The value of *H*_*U*_(*x, y*) or *H*_*L*_(*x, y*) at any point (*x, y*) was defined as the distance from the upper boundary of the 3D image to the upper or lower surface of the brain slice, respectively. Numerically, H_U_ and H_L_ were determined based on the optimization of 3 factors: (1) the value of the z-gradient image, which was generated by convolving the stitched image with a z-gradient filter; (2) the Laplacian of *H*_*U*_ and *H*_*L*_ as a smooth penalty of surface; (3) the distance between the upper and lower surfaces compared to the thickness of the slice (i.e. 300 μm physical distance). With these constraints, in the areas surrounding cortical sulci and ventricles, the boundaries of brain slices detected by the gradient filter which were not real physical cuts were not recognized as slice surfaces. The stitched slice image was then digitally resliced and flattened by moving and scaling along the Z-direction for each pixel, assuming both the upper and lower surfaces were horizontal, and the distance between any pair of *H*_*U*_(*x, y*) and *H*_*L*_(*x, y*) was 300 μm. The sets of *H*_*U*_ and *H*_*L*_ for all slices were used for further inter-slice stitching and SPS positioning.

#### Inter-slice stitching

Adjacent slices were stitched together after flattening by registering the upper surfaces of the *n*-th slice and the lower surfaces of the (*n-1*)-th slice. The registration was performed by *elastix*, using metrics of *mutual information* and *rigidity penalty*. The deformation field of surfaces was globally optimized using the stochastic gradient descent (SGD) algorithm, by minimizing the average deformation of the surface image and the average displacement between the upper and lower surfaces of a slice. Image volumes of each brain slice were transformed according to the resultant deformation field and then stacked into the whole brain image volume. Errors of this interslice stitching method were evaluated by manually recognizing 500 random pairs of axonal segments cut by the vibroslicer, in the reconstructed image volume of the brain hemisphere ipsilaterally injected with AAV-GFP into the MD, and calculating the average shift between the ends of fibers crossing those stitched surfaces.

#### Visualization

The whole monkey brain image was reconstructed at 10×10×10 μm^3^ voxel resolution. The reconstruction software also supports reconstructing ROIs of user-specified locations and sizes at full resolution. The image volumes of the whole brain or a given ROI were converted to the Imaris file format (IMS) for visualization in Imaris (v9.1∼9.5, Oxford Instruments) or in our custom software, Lychnis (see below). The IMS format is based on the standard hierarchical data format 5 (HDF5), which is open source and supports large image data. File format conversion was performed with the Imaris File Converter (v9.2, Oxford Instruments).

#### SMART positioning system (SPS)

A positioning system was established to bidirectionally map the pixels in the raw images to the reconstructed brain or ROIs. The positioning system contains three spaces: (1) Raw-image space *S*_*1*_(*s, k, x*_*r*_, *y*_*r*_, *z*_*r*_), where the arguments represent slice (*s*), stack (*k*), frame (*x*_*r*_), row (*y*_*r*_), and stack (*z*_*r*_) numbers of a pixel in *S*_*1*_; (2) Intra-slice-stitched-image space *S*_*2*_(*s, x*_*i*_, *y*_*i*_, *z*_*i*_) where the arguments represent the 3D position of a point (*x*_*i*_, *y*_*i*_, *z*_*i*_) at a stitched 3D slice (*s*) image; (3) Reconstructed-volume space *S*_*3*_(*x*_*v*_, *y*_*v*_, *z*_*v*_). The coordinate mapping between spaces are based on bidirectional transformations across the three spaces: (1) Resampling transformation between *S*_*1*_ and *S*_*2*_, consisting of a set of affine transformations each applied to an image stack; (2) Reconstruction transformation between *S*_*2*_ and *S*_*3*_, consisting of a group of displacement fields each applied for a slice image. This system is also extensible to the atlas space when necessary.

### Mesoscopic analysis

#### Fiber orientation analysis

We visualized fiber orientation by applying structure tensor analysis ^45^ on a low-resolution (10×10×10 μm^3^ voxel resolution) reconstruction of the whole brain volume. The structure tensor is defined as:

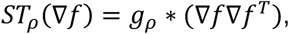

where ∇*f* is the gradient of the intensity *f*(*x, y, z*), of the reconstructed 3D image. *g*_ρ_ is a 3D Gaussian kernel with standard deviation ρ, set as 1 pixel here.

Fiber orientation *f_vis_* was given by the product of the secondary eigenvalue λ_2_ and eigenvector **v**_2_ of the structure tensor *ST*_ρ_(∇*f*):

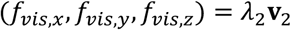

The resulted fiber orientation image was rendered and visualized in Imaris using the *blend* mode. The *x, y*, and *z* components of this fiber orientation image were rendered using red, green, and blue colors, respectively.

#### Cortical flattening

Surfaces of pial/gray (*pial*) and gray/white (*white*) boundaries were reconstructed based on structural MRI images (T1 MPRAGE, 250×250×500 µm^3^) of the same animal using Freesurfer ^46^ and custom Matlab (Mathworks) scripts. The mid surface between *pial* and *white* surfaces was inflated, and then 4 cuts were made to flatten the surface for visualization purposes (Supplementary Fig. 10). Cortical thickness was estimated by comparing *pial* and *white* surfaces. Atlas areas ^47^ were labeled onto the flattened surface based on nonlinear co-registration between the MRI of the test animal and the MRI template of the atlas, which were both nonlinearly warped to the National Institute of Mental Health Macaque Template ^48, 49^, serving as a common template. MD projection areas were determined by manually registering the coronal sections from the reconstructed whole-brain image to the atlas.

#### Parcellation

Cortical area identification was based on manually matching the DAPI, NT640 or autofluorescence images of the slices to the atlas ^47^ in the MIP images of each slice. Cortical layers were identified from the Nissl images or cellular autofluorescence in eGFP or mCherry channels.

### Fiber tracing

We developed a software referred to as “Lychnis” for tracing axonal fibers in a 3D image block generated by multiple imaging modules, or in the whole image set generated by the SMART pipeline, in which the users can mark nodes along the axonal tracks semi-automatically.

#### Fiber tracing in volume blocks

A reconstructed image volume at specified ROIs generated by the reconstruction software was converted to IMS format. Using high-level HDF5 API, small image blocks of a given size, e.g. 256×256×256 pixels at user-specified resolution and location, can be loaded from the IMS file in Lychnis, taking advantage of the multi-resolution structure and chunk-wise layout of HDF5. The Visualization Toolkit (VTK) ^50^ and Virtual Finger technology ^51, 52^ are used in Lychnis for 3D rendering and interactive labeling. Semi-automatic tracing was implemented by annotating two nearby markers in the axon and the tracks were automatically extended by linear extrapolation and manual correction when necessary. The nearby image blocks were automatically loaded for continuous tracing.

When necessary, misalignment that occurred during whole-brain reconstruction between adjacent slices resulting from the imperfect automatic volume reconstruction was manually corrected by virtue of the continuity of neuronal fibers and blood vessels in Lychnis. This software provides a user interface showing 3D visualization of two adjacent slice volumes, implemented with VTK, for the users to interactively mark the breakpoints of fibers on the surfaces of both slices. Then the deformation fields of all slices could be calculated based on these markers, using the interpolation library nn ^53^. Lychnis also provides bidirectional transformation between the image coordinate systems before and after deformation that was also integrated in the SMART positioning system.

#### Fiber tracing in whole macaque brains

Brain-wide axonal tracing was performed in the raw-image space with an adaptive and progressive approach implemented in Lychnis. All of the raw image stacks of the whole macaque brain were stored in OME-TIFF format at a remote data center. Lychnis provided a dynamic mechanism for loading and displaying the raw data while tracing. The starting axonal points or segments were selected in the low-resolution whole brain, and their locations were mapped to the raw-image coordinate system *S*_*1*_ via SPS. A block of raw images centering these points was loaded and assembled in memory by shifting individual images and rotating this local image block for 3D visualization in the Lychnis user interface. Tracing in this image block could be performed by the same semi-automatic algorithm for tracing in volumes of interests. When the fiber segment in the current slice was fully traced, the SPS converted the position of the last annotated point to a location in the whole-brain coordinate space *S*_*3*_, (*x*_*n*_, *y*_*n*_, *z*_*n*_), and then converted the nearby point (*x*_*n*_, *y*_*n*_, *z*_*n*_*+ε*) back to the raw-image space and loaded the corresponding image block, where *ε* was set to make sure *z*_*n*_ and *z*_*n*_+*ε* located in two neighboring slices. The fibers were traced with this iterative and progressive strategy block-by-block from the initial starting point or point set both anterogradely along the axon tracks to their endings in all branches, or retrogradely to the virus injection sites. In this study, axons from the injection site in the left hemisphere labeling MD and neighboring RE areas sparsely projecting to the contralateral cortical areas were traced. A few axons with significantly higher brightness could also be identified and traced out from dense ipsilateral fiber bundles. After fiber tracing was completed, all the points in the tracks were converted to the whole-brain space *S*_*3*_ for visualization. Completed tracing results were reviewed by two independent annotators, and the consensus axon tracks were used for the analysis ^19, 20^.

#### 3D rendering

A GPU-accelerated renderer was created and integrated into Lychnis for 3D rendering and creating videos. Imaris was also used for 3D rendering and creating videos.

## Supporting information

Supplementary materials (including Methods, Supplementary Figures and Tables)

## Acknowledgments

We thank Y. Song, M. Zhang, S. Zhao, T. Wang for their technical assistance with sample preparation and imaging. We thank Dr. Y. Guo and K. Zhang for viral validation in mice. We thank Dr. Yanyang Xiao, Dr. Shuo Chen, Dr. Pengcheng Zhou and Dasheng Bi for valuable suggestions on the manuscript. This work was supported by the Strategic Priority Research Program of Chinese Academy of Science (XDB32030200), the National Natural Science Foundation of China (91732304), the Key-Area Research and Development Program of Guangdong Province (2018B030331001; 2018B030338001), Shenzhen Infrastructure for Brain Analysis and Modeling (ZDKJ20190204002), and by Chinese Academy of Sciences International Partnership Program (172644KYSB20170004). Q.Z., L.I.Z., H.D., P.-M. L. and G.-Q.B were also partially supported by the NIH BICCN program (U01MH116990).

## Author contributions

Fang Xu, L.I.Z., H.D., Fuqiang Xu, X.H., P.-M. L. and G.-Q.B. conceptualize the project. Fang Xu led this project under supervision of P.-M. L. and G.-Q.B.. Fang Xu, Y.S. and H.W. established the pipeline for whole macaque brain imaging. Fang Xu, Q.Z., H.W. and C.X. designed and set up the microscope. Y.S. performed sample preparation and acquired data. L.D. developed the software for image acquisition, visualization, and neuronal tracing. C.Y. developed the software for brain reconstruction. H.T. and X.H. injected viruses and prepared the macaques brain samples. Fang Xu, Y.S., C.Y., F.W., and R.X. analyzed the data. Q.L. developed tools for image preprocessing. P.S. and Fuqiang Xu validated and provided the tracing viruses. H.D. and R.D. provided valuable neuroanatomical insights. Fang Xu, P.-M. L., and G.-Q.B. wrote the manuscript with inputs from all the authors.

## Data and materials availability

All codes used in this study are provided upon request. All data required to evaluate the conclusions in the paper are present in the paper or the supplementary materials. The complete imaging datasets of macaque brains exceed 1 petabyte, and therefore impractical to upload to a public data repository. The subsets related to any figure or video in this work are freely available upon request and providing feasible data transfer mechanisms (such as physical hard disk drives, cloud storage, or onsite visiting).

