## Supplementary materials (including Methods, Supplementary Figures and Tables) for "High-throughput whole-brain mapping of rhesus monkey at micron resolution"

### Supplementary Figures, Tables and Video legends

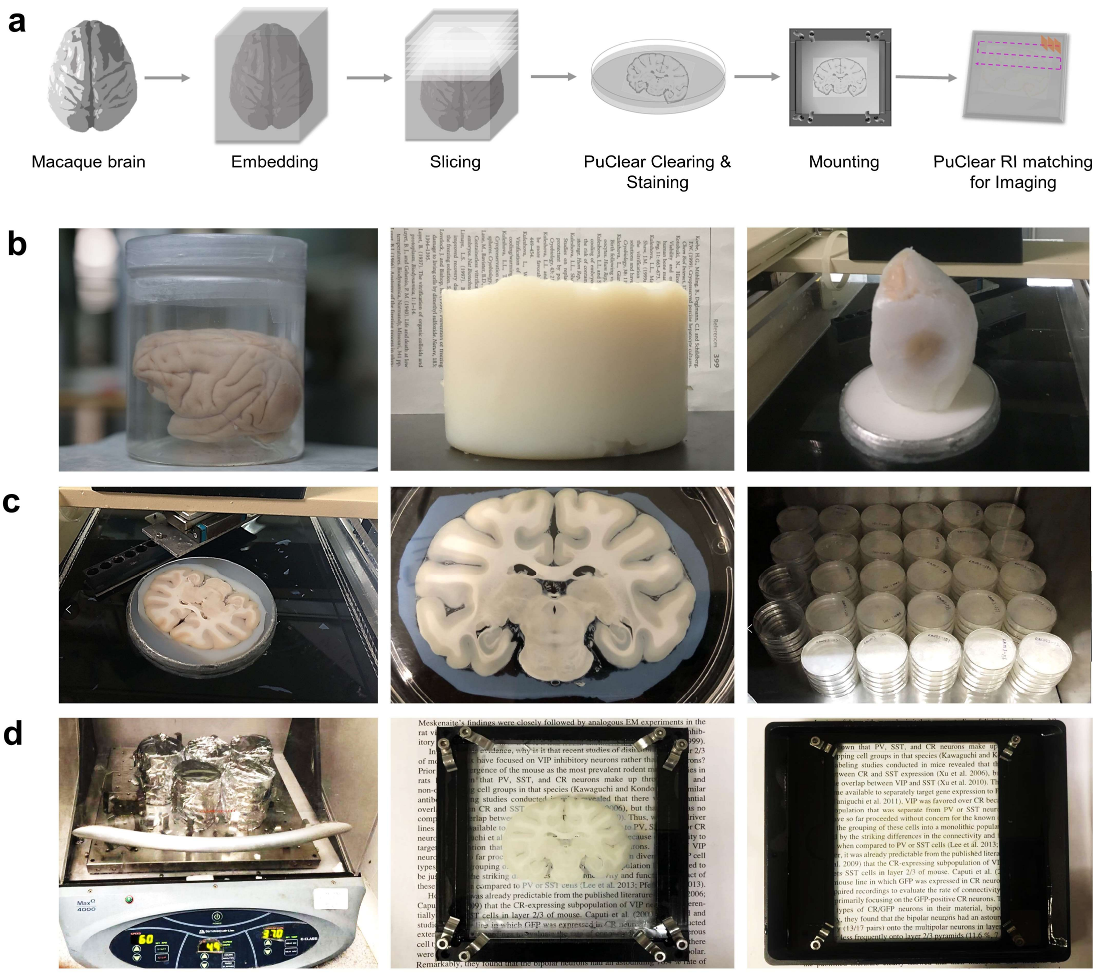


**Supplementary Fig. 1 | The SMART sample preparation workflow.** (**a**) Key steps of sample preparation workflow. (**b**) Embedding. The excised monkey brain was post-fixed with 4% HMS (*left*) and embedded at 37°C until polymerization into a tissue block (*middle*). The block was trimmed to remove extra polymer surrounding the brain tissue (*right*). (**c**) Slicing. The embedded brain was constrained in a tube in the compresstome slicer, with the extra space in the tube filled with low melt agarose. After the agarose congealed, the brain was sectioned into 300-μm-thick slices (*left*), each transferred into a Petri dish with 40 mL PBS (*middle*). In total, about 250 slices of each entire brain were collected and then stored in a 4°C fridge before clearing (*right*). (**d**) PuClear clearing and mounting. The slices were treated with PuClear clearing solution (5% PBST) to increase cellular permeability at 37°C with gentle shaking (*left*), then mounted on a glass substrate by virtue of a custom chamber (*middle*) after optional staining. Brain slices attached to glass substrates were incubated in the PuClear RI matching solution for 1 h in the imaging chamber until transparent (*right*).

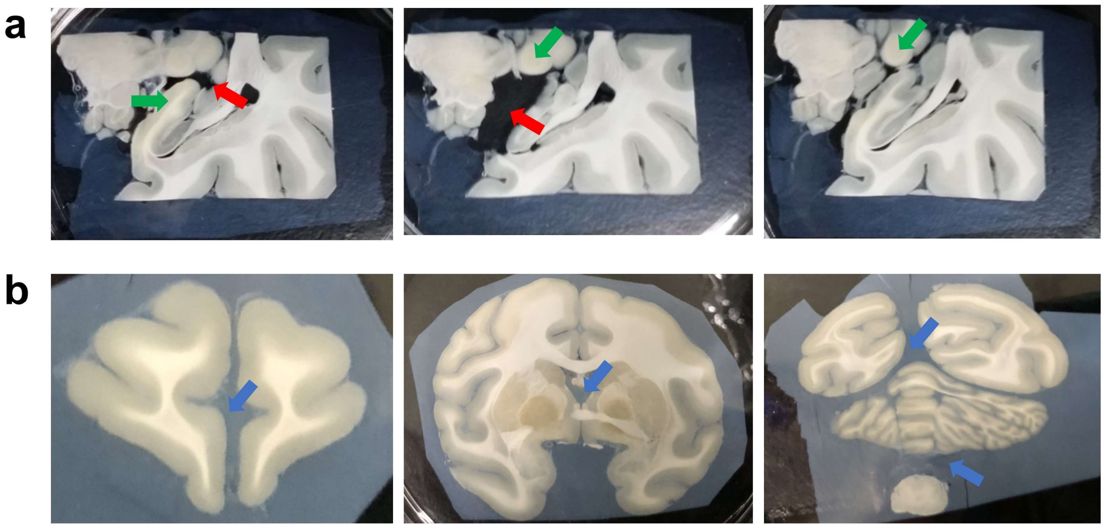

**Supplementary Fig. 2 | Hydrogel-based embedding preserves slice structures.** (**a**) Consecutive slices sectioned after normal agarose embedding of monkey brain fixed by PFA. Arrows indicate tissue loss (red) and non-uniform slice thickness (green). (**b**) Slices of a macaque brain sectioned after hydrogel-based embedding. Ventricles and the space between disconnected parts of the brain were filled with the crosslinked hydrogel (blue), which preserved the morphological integrity of slices.

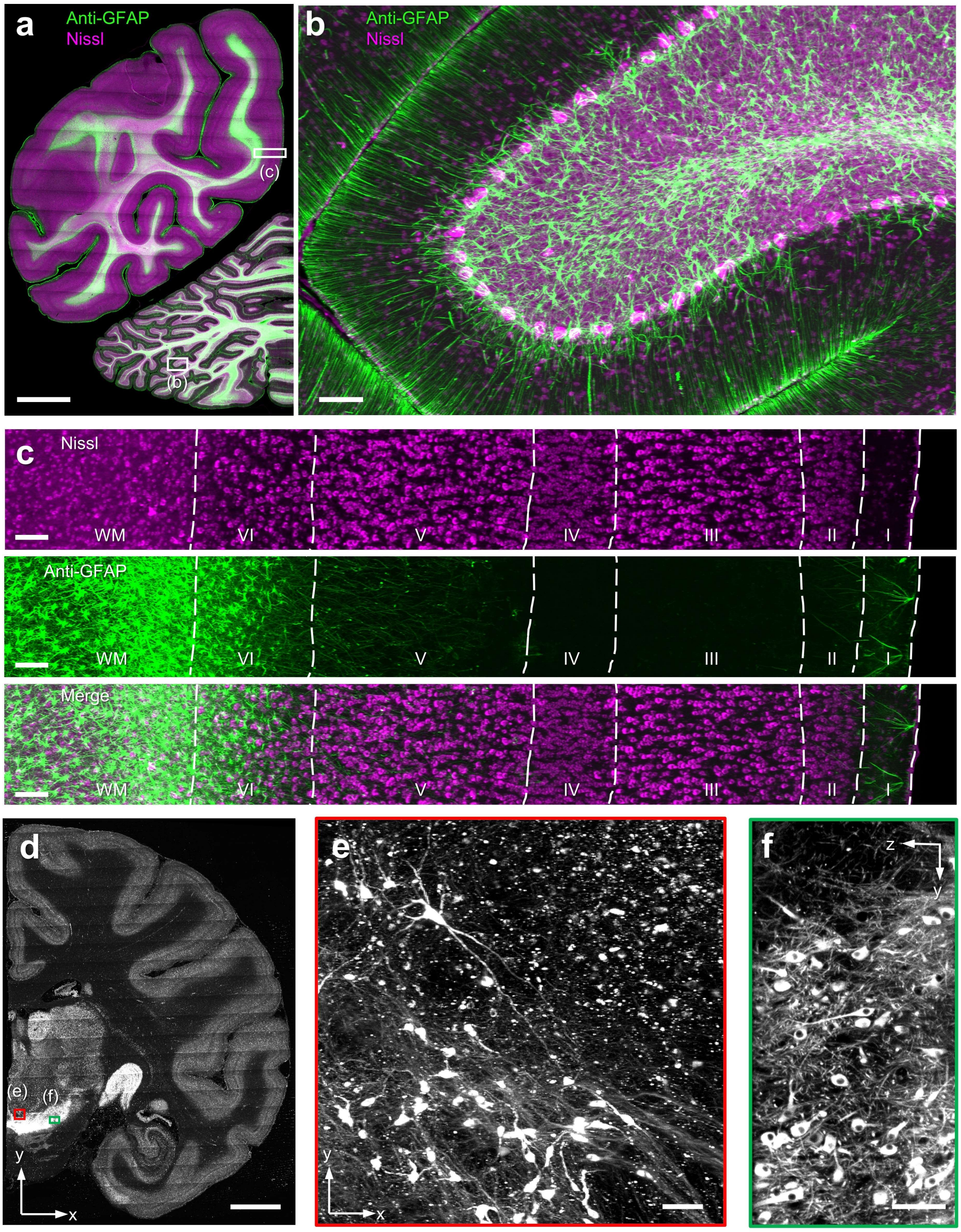


**Supplementary Fig. 3 | PuClear is compatible with immunolabeling.** (**a**) Maximum intensity projection (MIP) of a slice with neurons labeled with NT640 for Nissl staining, and with anti-glial fibrillary acidic protein (GFAP) antibodies for astrocyte labeling. Images were acquired with the VISoR2 microscope after PuClear treatment. Boxed ROIs are enlarged and shown in (b) and (c). (**b**) An ROI in the cerebellum showing the characteristic Purkinje cells and astrocyte fibers. (**c**) An ROI in the cerebral cortex showing the laminar distribution of neurons and astrocytes. Cortical layers were recognized by Nissl patterns. (**d-f**) MIP of a slice stained with anti-tyrosine hydroxylase (TH) antibodies for labeling dopamine neurons (d). Two ROIs in the 3D image of substantia nigra (SN) were magnified and shown in (e) (X-Y view) and (f) (Y-Z view). Scale bars: (a) and (d), 5 mm; (b), (c), (e) and (f), 100 μm.

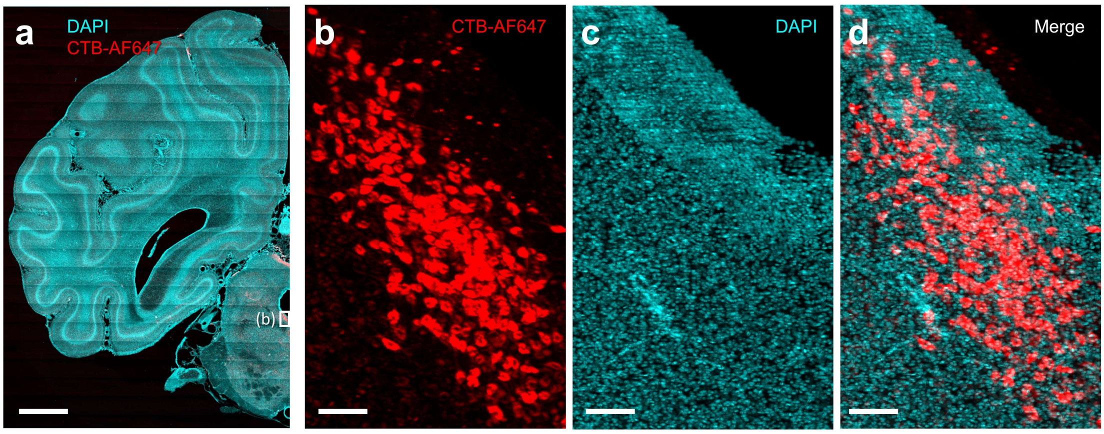


**Supplementary Fig. 4 | PuClear is compatible with CTB retrograde labeling.** (**a**) MIP of a slice from a brain labeled with CTB-AF647 for 2 months and post-stained with DAPI after PuClear clearing. (**b-d**) Magnified views of the box region in (a), showing the CTB-labeled neurons. Scale bars: (a), 5 mm; (b), (c) and (d), 100 μm.

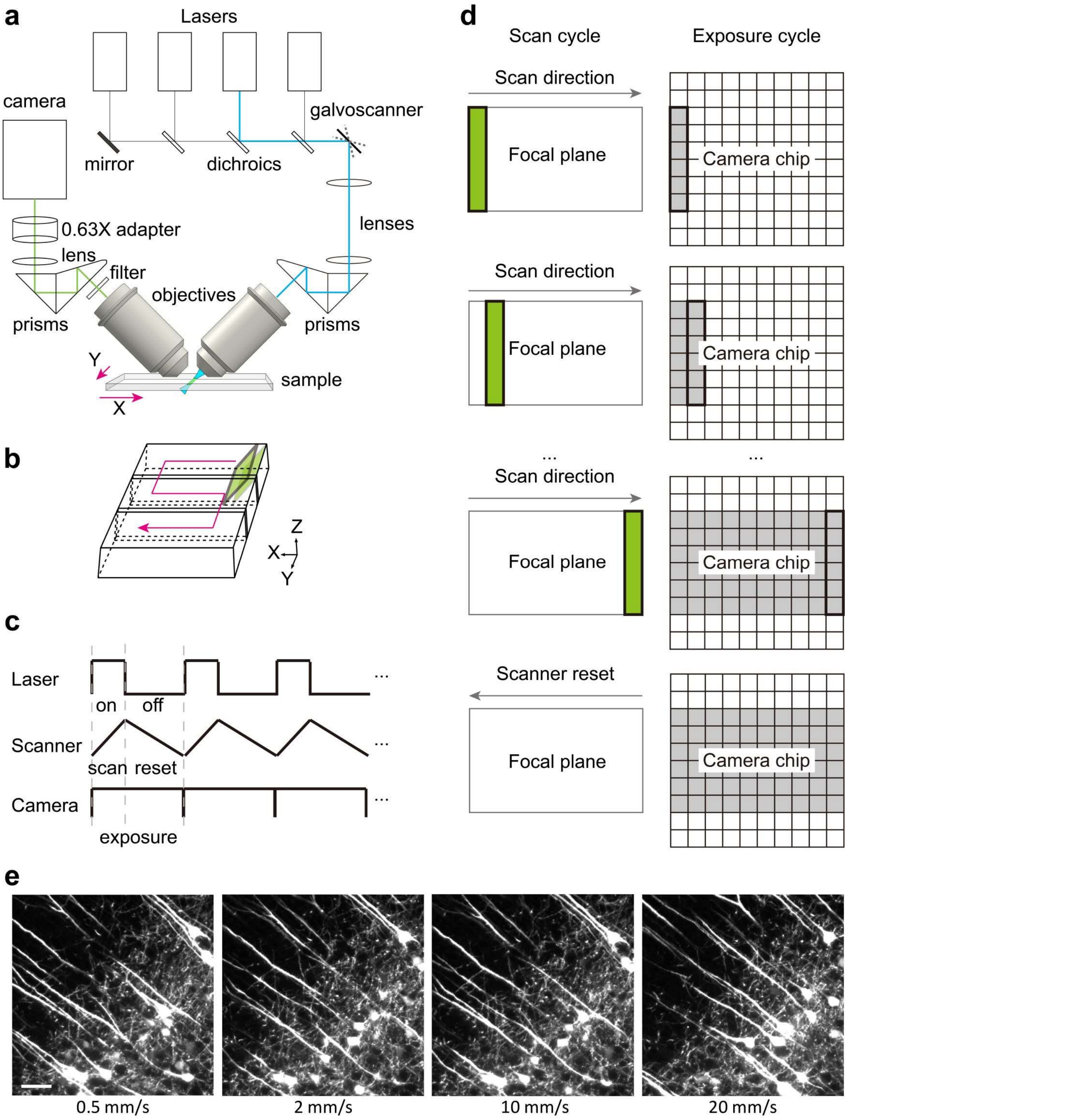


**Supplementary Fig. 5 | Principles of high-throughput VISoR2 microscopy.** (**a**) The schematic light path of the VISoR2 microscope. The illumination (right) and imaging (left) objectives are both 45° oblique to the sample surface. (**b**) Motion control of VISoR2. The sample was moved continuously in an "S" style while images were acquired. (**c**) Sequence control of VISoR2. (**d**) Illustration depicting the synchronization of the scan cycle and imaging cycle, which erases motion blurs. (**e**) Example images of a Thy1-transgenic mouse brain acquired while the brain slice moved at different velocities along the X-axis. No motion blurring was noticed. Scale bar: 50 μm.

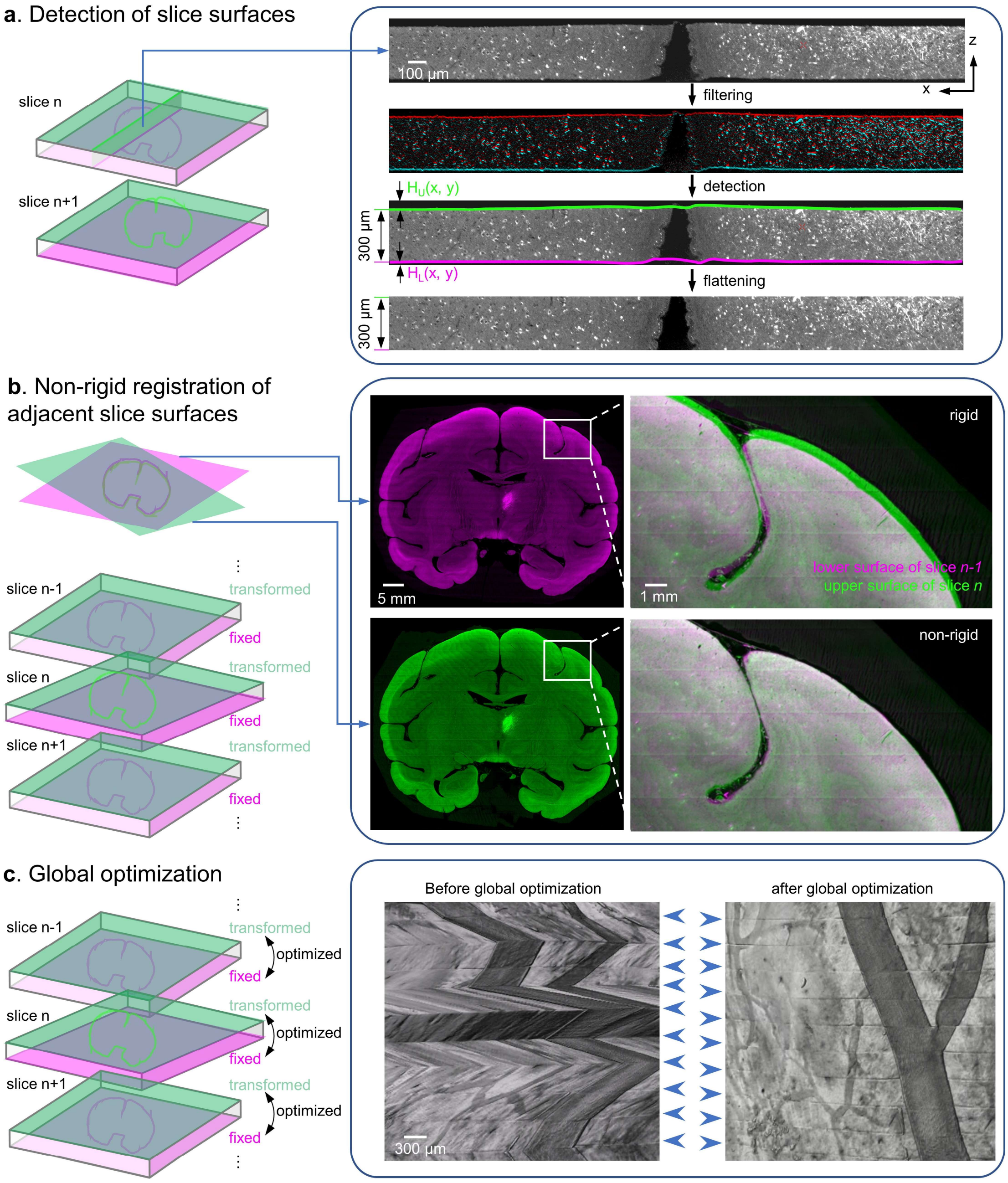


**Supplementary Fig. 6 | The volume stitching pipeline for whole-brain reconstruction.** (**a**) Steps of slice flattening. A slice image was Z-gradient filtered, and the surfaces were detected and optimized iteratively. The 3D slice image was then flattened based on the physical distance (i.e. 300 µm) between surfaces. (**b**) Initial inter-slice registration was performed by non-rigidly aligning the upper surface of a slice to the lower surface of its precedent slice. (**c**) Additional global optimization was applied to finely tune the inter-slice registration by minimizing the intra-slice deformation. Arrowheads indicate the stitches between slices.

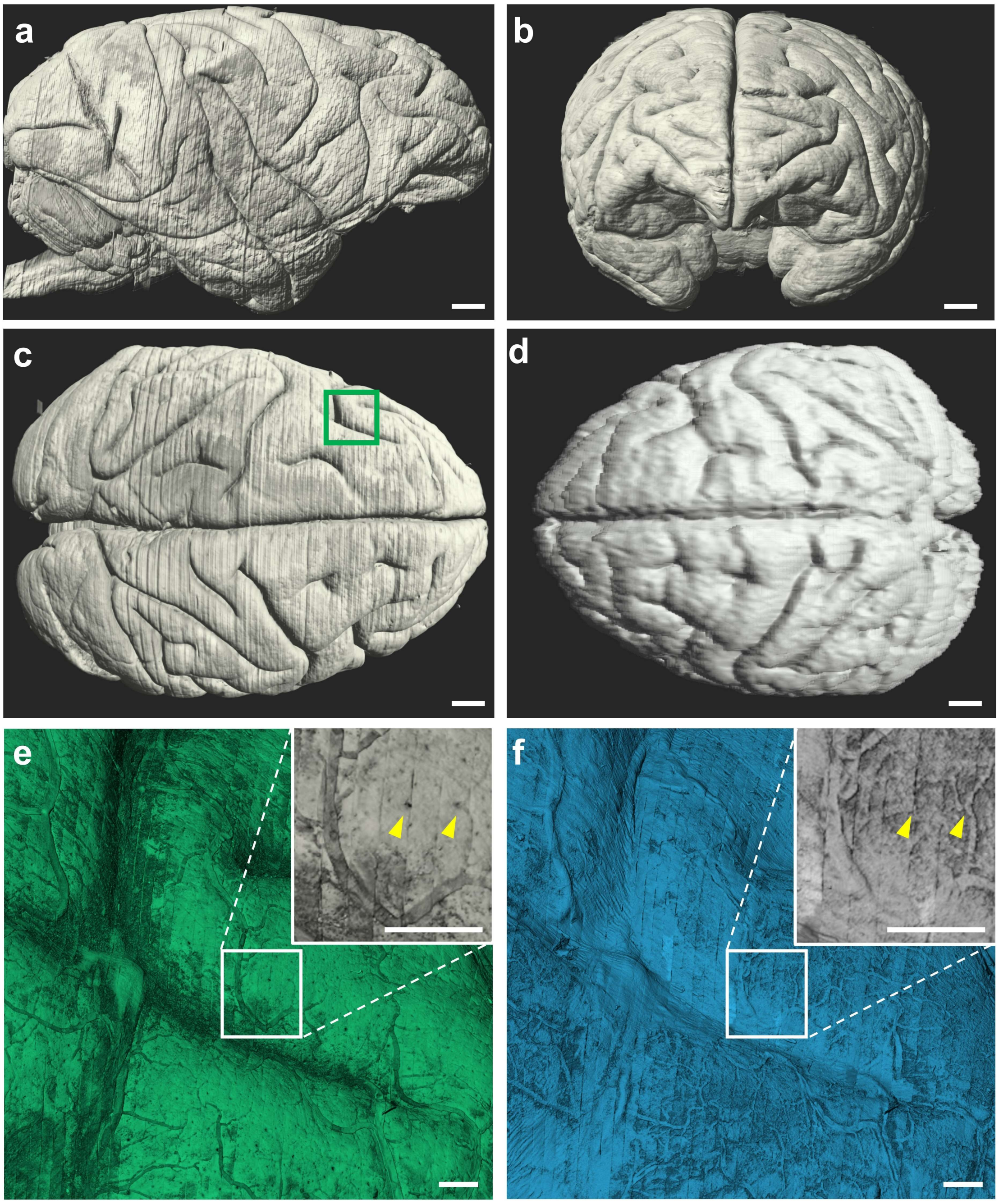


**Supplementary Fig. 7 | Reconstructed whole macaque brain.** (**a-c**) A macaque brain reconstructed from its complete set of slices imaged with VISoR2. Three perspectives visualized from the right (a), front (b), and top (c) sides are shown, respectively. Brain autofluorescence was rendered in the *normal shading* mode in Imaris. Scale bars: 5 mm. (**d**) MRI image of the same brain acquired at 0.25×0.25×0.5 mm$^3$ resolution. Scale bar, 5 mm. (**e-f**) Magnified view of the boxed area in (c) shows the detailed vascular structures the brain surface imaged for 488 nm-excited autofluorescence (e) or DAPI fluorescence (f). Densely distributed tiny pores formed by the vessel penetration (examples indicated by yellow arrowheads) were observed. Scale bars: 1 mm.

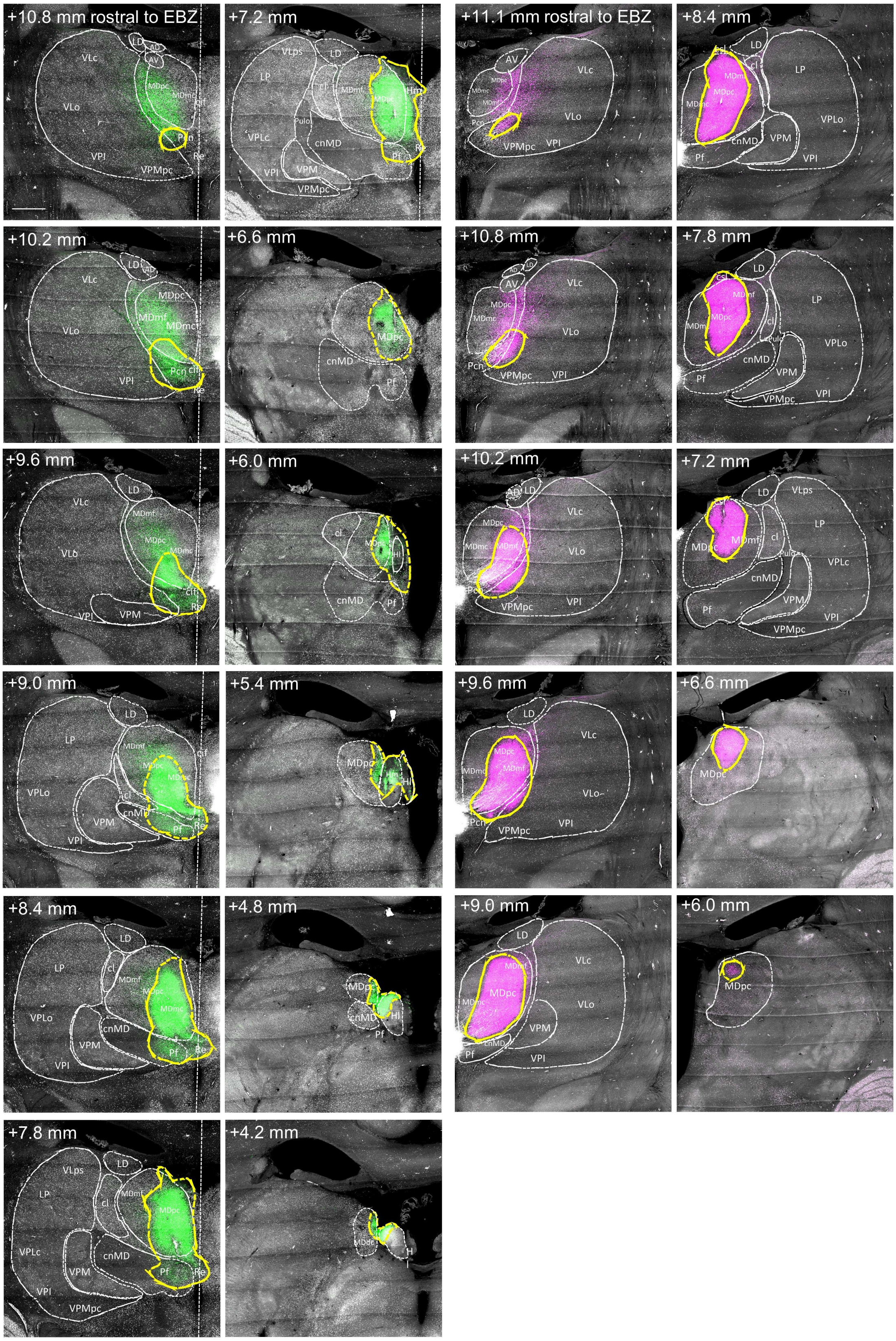


**Supplementary Fig. 8 | Sample images of the AAV injection sites.** Sections through the AAV-eGFP injection sites (left two columns) and those through the AAV-mCherry injections sites (right two columns) are shown, respectively. Brain areas were annotated with dashed lines in white. Areas with neuron somata were outlined with dashed lines in yellow. Straight dashed lines indicate the midline. Scale bar: 2 mm. Acronyms: AV, anterior ventral nucleus; cif, central inferior nucleus; cl, central lateral nucleus; cnMD, centromedian nucleus; csl, central superior lateral nucleus; Hl, lateral habenular nucleus; LD, lateral dorsal nucleus; LP, lateral posterior nucleus ; MDdc, medial dorsal nucleus, densocellular division; MDmc, medial dorsal nucleus, magnocellular division; MDmf, medial dorsal nucleus, multiform division; MDpc, medial dorsal nucleus, parvicelluar division; Pcn, paracentral nucleus; Pf, parafascicular nucleus; Pulo, pulvinar oralis nucleus; Re, reunions nucleus; VLc, ventral lateral caudal nucleus; VLo, ventral lateral oral nucleus; VLps, ventral lateral postrema nucleus; VPI, ventral posterior inferior nucleus; VPLc, ventral posterior lateral caudal nucleus; VPLo, ventral posterior lateral oral nucleus; VPM, ventral posterior medial nucleus; VPMpc, ventral posterior medial nucleus, parvicellular division.

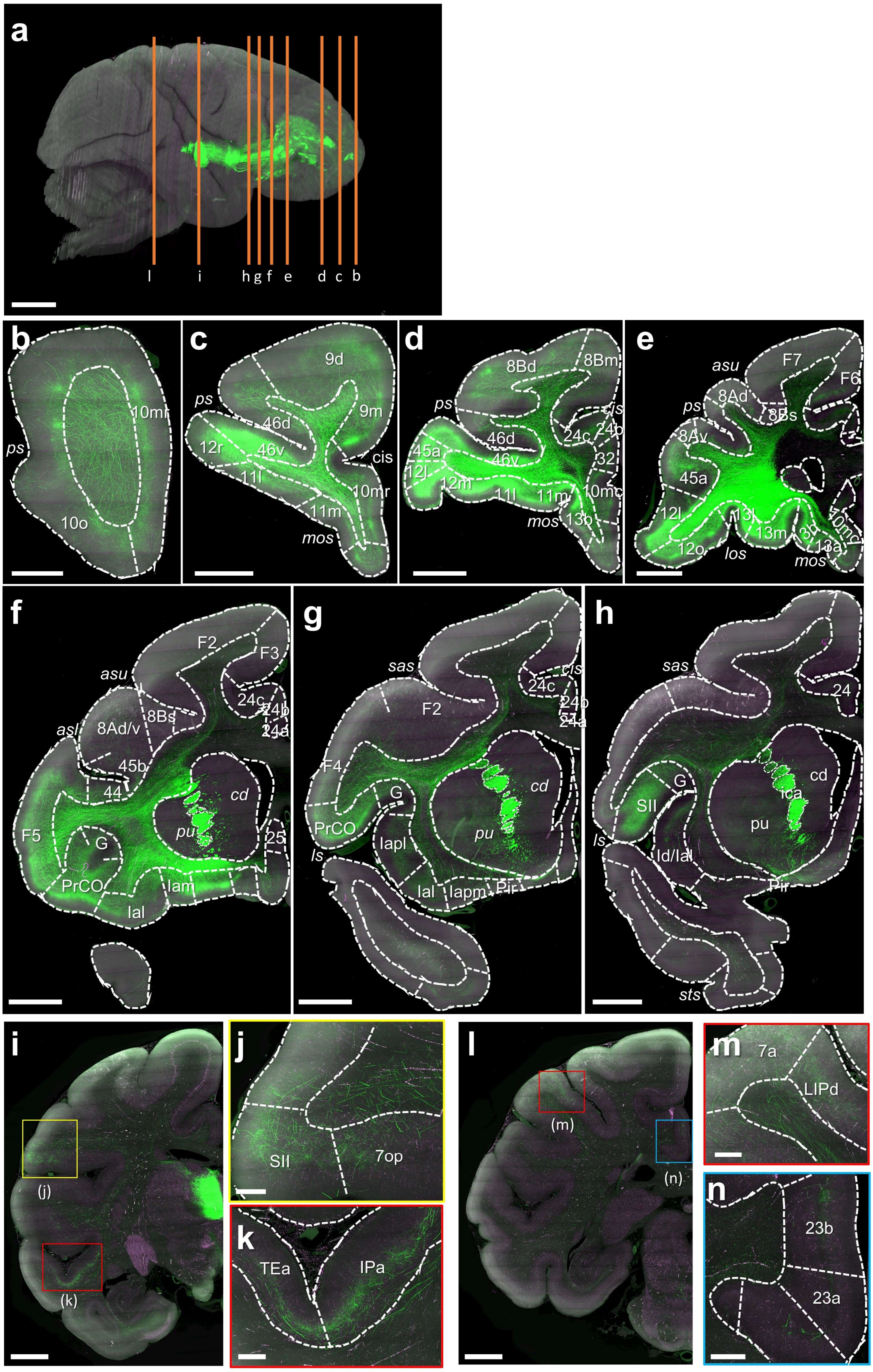


**Supplementary Fig. 9 | Thalamocortical projection of MD.** (**a**) Side view of the reconstructed monkey brain. Only the left hemisphere injected with AAVs in MD areas are shown. Vertical lines indicate the location of the example coronal sections shown in (**b-i**) and (**l**). (**j-k**) Enlargement views of boxed areas in (**i**). (**m-n**) Enlargement views of boxed areas in (l). Scale bars: (a), 10 mm; (b), 2 mm; (c-i), (l), 5 mm; (j-k), (m-n), 1 mm. Acronyms: *asl*, arcuate sulcus lower limb; *asu*, arcuate sulcus upper limb; cd, caudate nucleus; *cis*, cingulate sulcus; *ica*, internal capsule anterior limb; *los*, lateral orbital sulcus; *ls*, lateral sulcus; *mos*, medial orbital sulcus (and branch); *ps*, principal sulcus; pu, putamen; *sas*, spur of the arcuate sulcus; *sts*, superior temporal sulcus; others are listed in Supplementary Table s1.

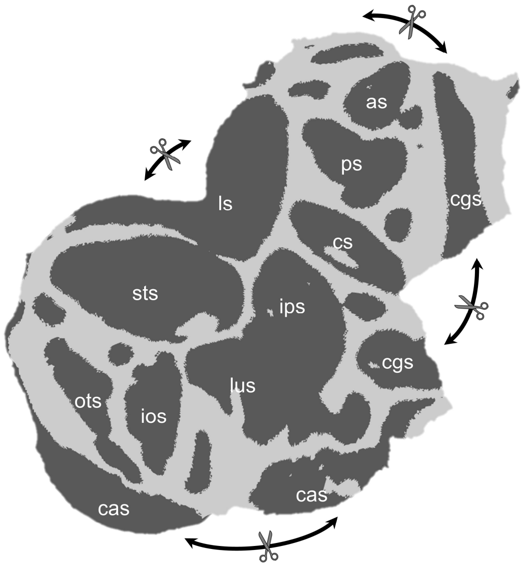


**Supplementary Fig. 10 | Cortical flattening.** Four cuts were made at specified locations to flatten the cortical surface. Acronyms: *as*, arcuate sulcus; *cs*, central sulcus; *cas*, calcarine sulcus; *cgs*, cingulate sulcus; *io*, inferior occipital sulcus; *ips*, intraparietal sulcus; *ls*, lateral sulcus; *lus*, lunate sulcus; *ots*, occipitotemporal sulcus; *ps*, principal sulcus; *sts*, superior temporal sulcus.

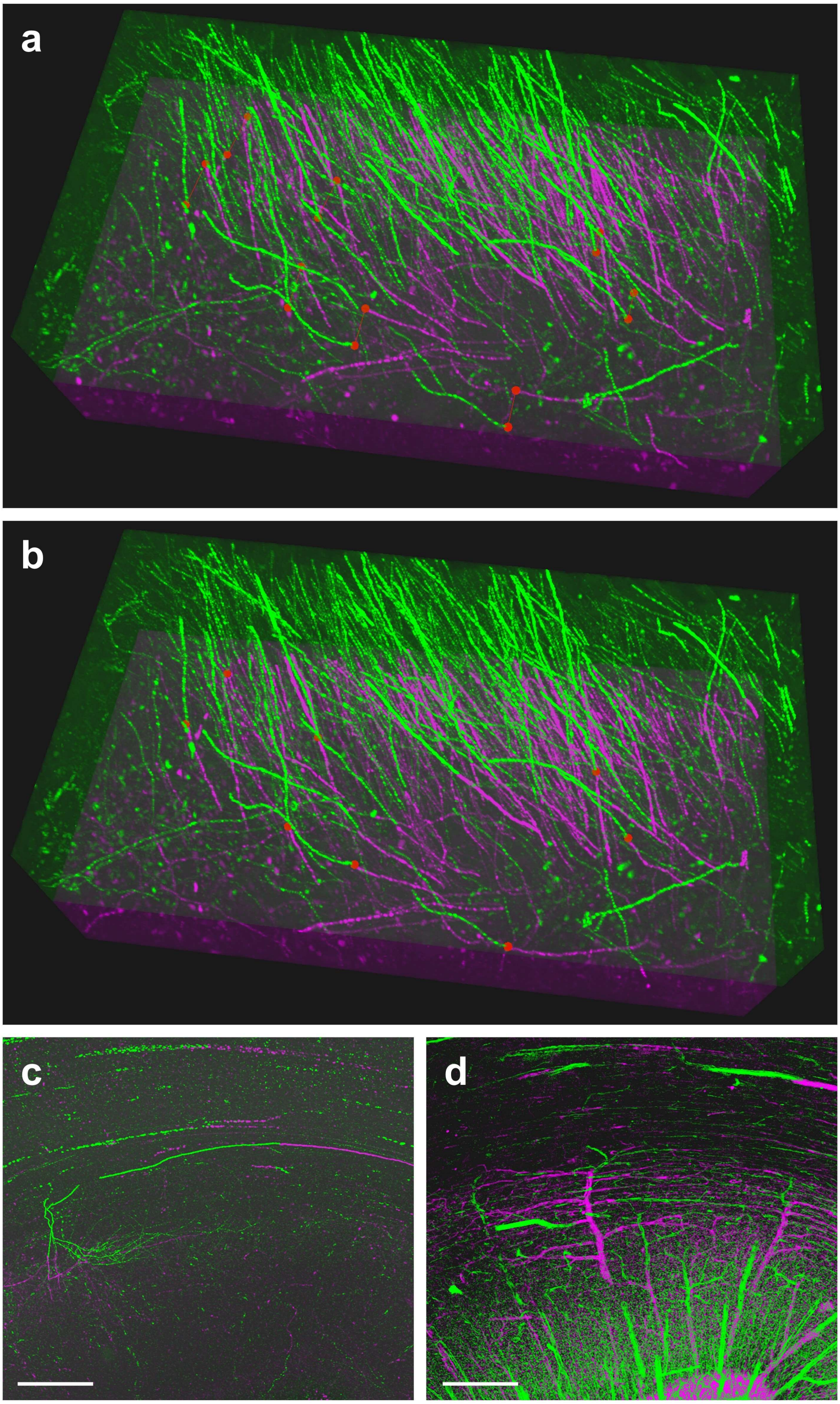


**Supplementary Fig. 11 | Correction of slice misalignment.** (**a**) A snapshot showing two adjacent slices (separately colored green or magenta) 3D rendered in Lychnis stitcher which allows users to identify and annotate a set of paired endpoints (red) of axonal fibers. (**b**) A snapshot showing the slices in (a) after alignment according to an XY deformation field that was generated from the annotations. (**c-d**) In areas with sparse axons (c), dense vascular structures of the same ROI are visible by reversing the image look-up tables (d). The lumen of blood vessels are filled with HMS during perfusion which is non-fluorescent and thus darker than the brain tissues in the images (c). In this reversed rendering mode provided in Lychnis, the vessels are revealed as a reference dense fibrous framework that assists refining misalignment. Scale bars: 500 μm.

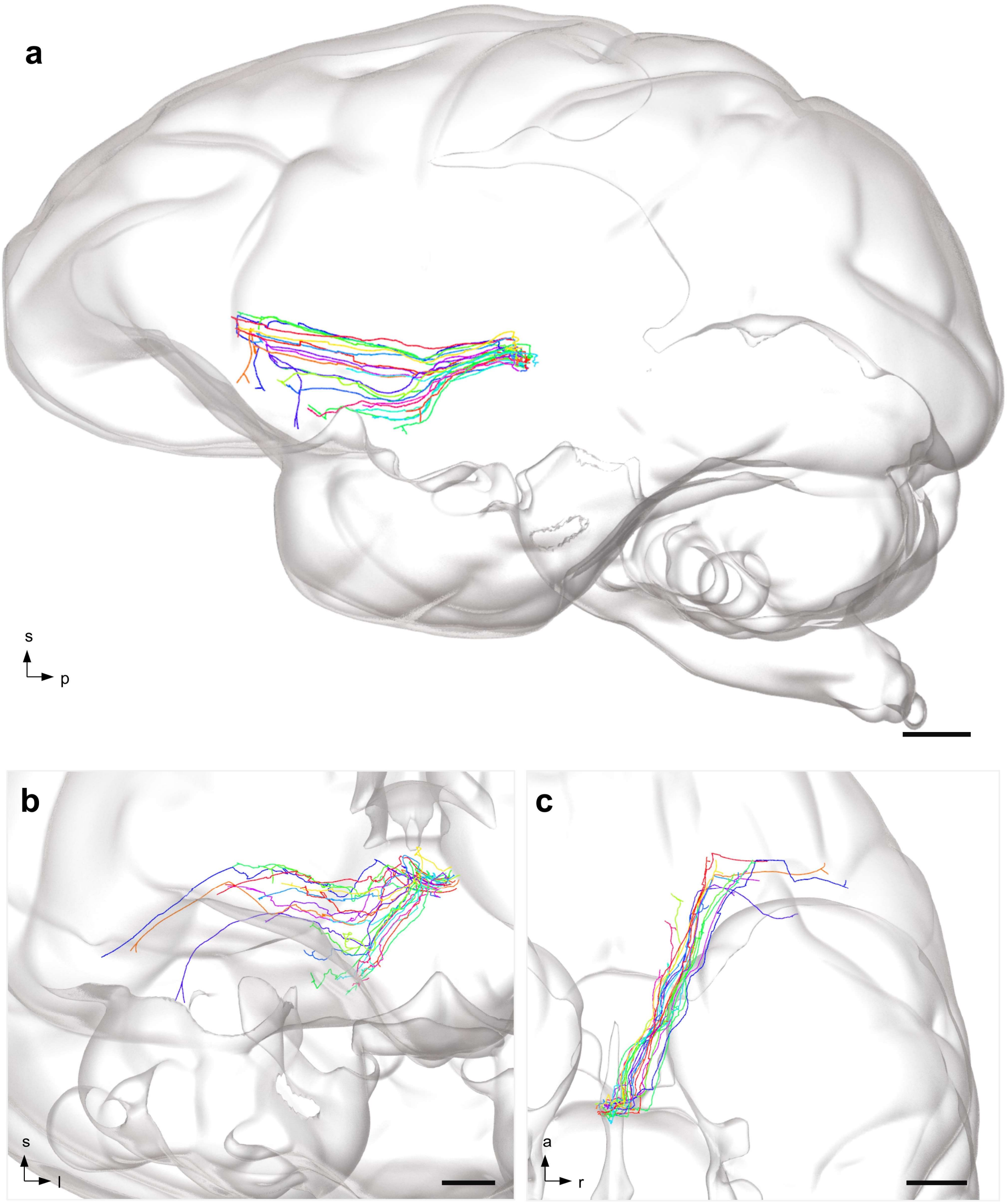


**Supplementary Fig. 12 | Fiber segments in the internal capsule.** Twenty-two fibers traveling in the internal capsule were traced and the segments from the MD and RE areas to their first branching points are shown in side (**a**), front (**b**) and top (**c**) views. Scale bars: (a), 5 mm; (b), 3 mm; (c), 5 mm.

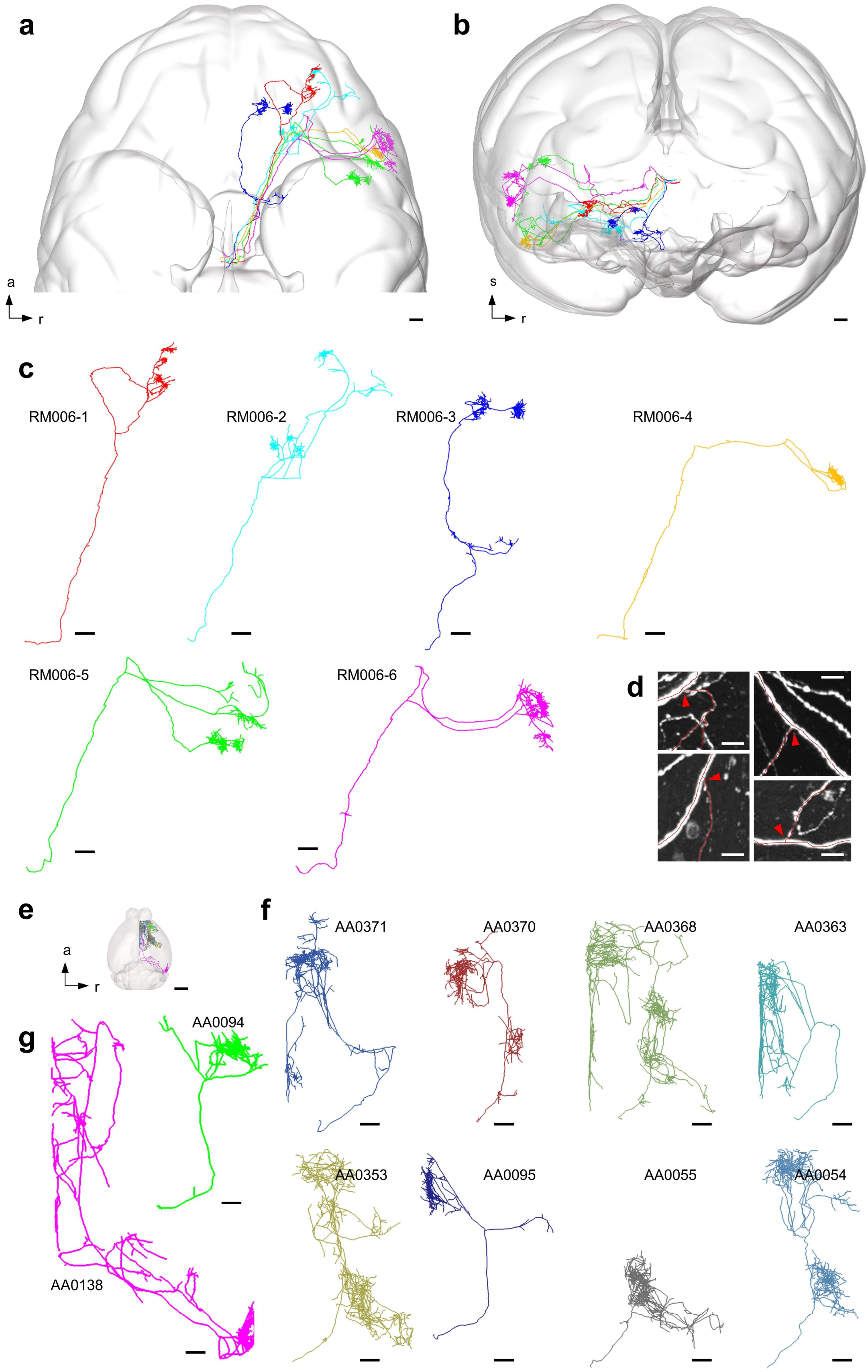


**Supplementary Fig. 13 | Comparison of fibers in macaque and mouse brains.** (**a-b**) Six macaque axons shown in top view (**a**) and front view (**b**), with the brain contours as a reference. Orientations: a, anterior; r, right; s, superior. Scale bars, 2 mm. (**c**) Morphology of individual fibers. Scale bars, 2 mm. (**d**) Example subcortical bifurcations in macaque neurons. Scale bars, 20 µm. (**e**) 8 mouse MD axons and 2 RE axons shown with the brain contour as a reference. Scale bar: 2 mm. (**f-g**) Enlarged views of each MD (AA0371, AA0370, AA0368, AA0363, AA0353, AA0095, AA0055, AA0054) and RE axons (AA0138, AA0094). Scale bars: 400 µm. Mouse neuron data are acquired from MouseLight project under CC-BY NC license [20, 54].

**Supplementary Table 1 | Summary of ipsilateral neuronal projections from the injection sites in the macaque brain.**

| Acronym | Full name of brain areas | Projection (eGFP) * | Projection (mCherry) | Reported in Ref (*26*) |
|---|---|---|---|---|
| 7a | Caudal inferior parietal lobule area (or Opt/PG) | low | | |
| 7b | Rostral inferior parietal lobule area (or PFG/PF) | low | | |
| 7op | Area 7op (parietal operculum) | low | | |
| 8Ad | Periarcuate area (or frontal eye field), dorsal subdivision | high | ** | # |
| 8Av | Periarcuate area (or frontal eye field), ventral subdivision | low | ** | # |
| 8Bs | Dorso-lateral prefrontal area in the arcuate sulcus upper limb | low | ** | # |
| 8Bd | Dorso-lateral prefrontal area | low | ** | # |
| 8Bm | Dorso-medial prefrontal area | low | ** | # |
| 9d | Dorso-lateral prefrontal area | high | ** | # |
| 9m | Dorso-medial prefrontal area | low | ** | # |
| 10mc | Medial prefrontal area | low | ** | |
| 10mr | Medial prefrontal area | high | ** | |
| 10o | Orbital prefrontal area | low | ** | |
| 11l | Orbital prefrontal area | high | ** | # |
| 11m | Orbital prefrontal area | high | ** | # |
| 12l | Ventro-lateral prefrontal area | high | ** | |
| 12m | Orbital prefrontal area | high | ** | |
| 12o | Orbital prefrontal area | high | ** | |
| 12r | Ventro-lateral prefrontal area | high | ** | |
| 13a | Orbital prefrontal area | low | | # |
| 13b | Orbital prefrontal area | high | ** | # |
| 13l | Orbital prefrontal area | high | ** | # |
| 13m | Orbital prefrontal area | high | ** | |
| 23a | Subregion of posterior cingulate cortex | low | | |
| 23b | Subregion of posterior cingulate cortex | low | | |
| 23c | Subregion of posterior cingulate cortex | low | | |
| 24a | Subregion of anterior cingulate cortex | low | | # |
| 24b | Subregion of anterior cingulate cortex | low | ** | # |
| 24c | Subregion of anterior cingulate cortex | low | ** | # |
| 25 | Medial prefrontal area (subgenual cortex) | low | | |
| 32 | Medial prefrontal area | high | ** | # |
| 44 | Ventro-lateral prefrontal area in the arcuate sulcus lower limb | high | ** | |
| 45a | Ventro-lateral prefrontal area | high | ** | # |
| 45b | Ventro-lateral prefrontal area in the arcuate sulcus lower limb | high | ** | # |
| 46d | Prefrontal area in the dorsal bank and lip of the principal sulcus | high | ** | # |
| 46f | Prefrontal area in the fundus of principal sulcus | high | ** | # |

| Acronym | Full name of brain areas | Projection (eGFP) * | Projection (mCherry) | Reported in Ref (*26*) |
|---|---|---|---|---|
| 46v | Prefrontal area in the ventral bank and lip of the principal sulcus | high | ** | # |
| F2 | Agranular frontal area F2 (or 6DR/6DC) | low | ** | |
| F3 | Agranular frontal area F3 (or SMA) | low | | |
| F4 | Agranular frontal area F4 (or 4C/6Va/6Vb) | low | ** | |
| F5 | Agranular frontal area F5 (or 6Va/6Vb) | high | ** | |
| F6 | Agranular frontal area F6 (or preSMA) | low | | |
| F7 | Agranular frontal area F7 (or 6DR) | low | | |
| G | Gustatory cortex | low | | |
| Iai | Intermediate agranular insula area | high | ** | # |
| Ial | Lateral agranular insula area | low | | # |
| Iam | Medial agranular insula area | high | ** | # |
| Iapl | Posterolateral agranular insula area | low | ** | # |
| Iapm | Posteromedial agranular insula area | high | ** | # |
| Id | Dysgranular insula | low | | |
| Ig | Granular insula | low | ** | |
| IPa | Sts fundus area | low | | |
| LIPd | Lateral intraparietal area, dorsal subdivision | low | ** | |
| PGa | Sts fundus/dorsal bank area | low | | |
| Pir | Piriform cortex | low | ** | |
| PrCO | Precentral opercular area | high | ** | |
| RM | Rostromedial, belt region of the auditory cortex | low | | |
| SII | Secondary somatosensory area | low | ** | |
| TAa | sts dorsal bank area | low | | |
| TEa | sts ventral bank area | low | ** | |
| TEm | sts ventral bank area | low | | |
| TEpv | Ventral subregion of posterior TE | low | | |
| TPO | sts dorsal bank area | low | | |
| Tpt | Temporo-parietal area | low | | |

*: Ipsilateral projection density is determined by thresholding the fluorescence intensity of the cortical flat map to reveal relative projection density from the AAV-eGFP injection sites in the left MD areas.

**: Cortical areas in the right hemisphere with mCherry-labeled fibers originated from right MD injection sites are noted.

#: Cortical areas that receive MD projections are noted based on ref. (*26*).

**Supplementary Table 2 | Comparison of macaque and mouse thalamic axonal projections.**

Mouse neuron data are acquired from MouseLight project under CC-BY NC license (*20*, *54*). Somata of neurons AA0054, AA0055, AA0095, AA0353, AA0363, AA0368, AA0370, AA0371 are located at the mediodorsal nucleus (MD). Somata of neurons AA0094 and AA0138 are located at the nucleus of reunions (RE).

Acronyms (macaque): same as in Supplementary Table 1.

Acronyms (mouse): ACAd, anterior cingulate area, dorsal part; ACAv, anterior cingulate area, ventral part; AId, agranular insular area, dorsal part; AIv, agranular insular area, ventral part; FRP, frontal pole; GU, gustatory areas; ILA, infralimbic area; MOp, primary motor area; MOs, secondary motor area; OLF, olfactory areas; ORBl, orbital area, lateral part; ORBm, orbital area, medial part; ORBvl, orbital area, ventrolateral part; PERI, perirhinal area; PIR, piriform area; PL, prelimbic area; RSPv, retrosplenial area, ventral part; VISpl, posterolateral visual area.

| **Macaque Axon ID (this study)** | **Cortical targets** | **Mouse Neuron ID (MouseLight data)** | **Cortical targets** |
|---|---|---|---|
| RM006-1 | 11l, 12m,12r | AA0054 | ACAd, AId, AIv, FRP, GU, MOp, MOs, ORBl, PL |
| RM006-2 | 11l, 12l, 12m, 12r, 13l, 13m | AA0055 | (none) |
| RM006-3 | 13b, 13m | AA0095 | AIv, FRP, ILA, ORBm, ORBvl, PL |
| RM006-4 | F5, PrCO | AA0353 | AIv, FRP, MOp, MOs, ORBl, ORBm, ORBvl, PL |
| RM006-5 | F5, PrCO, SII | AA0363 | ACAd, ACAv, ILA, Isocortex, MOp, MOs, PL, RSPv |
| RM006-6 | F4, F5 | AA0368 | ACAd, ACAv, AId, AIv, MOp, MOs, ORBl, ORBm, ORBvl, PL, RSPv |
| | | AA0370 | ACAd, ACAv, ILA, MOs, ORBl, ORBm, ORBvl, PL |
| | | AA0371 | ACAd, ACAv, FRP, MOp, MOs, OLF, ORBl, ORBm, ORBvl, PL, RSPv |
| | | AA0094 | AId, AIv, ILA, ORBl, ORBm, ORBvl, PIR |
| | | AA0138 | ACAd, ACAv, ILA, MOp, MOs, ORBl, ORBm, ORBvl, PERI, PL, RSPv, VISpl |

**Supplementary Video 1.**

On-the-fly VISoR2 imaging of a 300 µm-thick brain slice from a virus-injected adult macaque at 1 × 1 × 2.5 $\mu m^3$ resolution in 142 seconds. This video is at 2X playback speed. Fibers labeled with eGFP in the prefrontal area are revealed.

**Supplementary Video 2.**

A stitched image volume spanning 4 slices of a macaque brain.

**Supplementary Video 3.**

A reconstructed macaque brain. This volume was acquired at 1 × 1 × 2.5 $\mu m^3$ voxel resolution and the reconstructed volume was downsampled to 10 × 10 × 10 $\mu m^3$ resolution for rendering. A fiber orientation image of this brain is shown. Efferent fibers from the injection sites are labeled.

**Supplementary Video 4.**

Representative axons segmented from an ROI in the temporal lobe show distinct turning patterns.

**Supplementary Video 5.**

An axon traced from the injection site (covering MD and part of RE) to the contralateral cortical area F5 (the main segment of axon #RM006-4) and visualized in the whole-brain framework.
